# CsubMADS1 overexpression is associated with increased CAB transcript levels and with enhanced tolerance under nutrient limitation

**DOI:** 10.64898/2025.12.15.690641

**Authors:** Saraswati Nayar

## Abstract

MADS-box transcription factors (TFs) have been extensively studied in seed plants, but their functions in non-seed plants remain comparatively underexplored. CsubMADS1, a MADS-box TF from the microalga *Coccomyxa subellipsoidea* C-169, was previously shown to influence developmental processes and starvation-associated responses. In this study, overexpression of *CsubMADS1* was associated with increased tolerance to nutrient-limiting conditions, particularly during stationary phase and nitrogen starvation. Overexpressing lines exhibited reduced neutral lipid droplet formation and decreased mucilage staining under these conditions—physiological traits typically linked to stress-associated responses in microalgae. RNA-seq analysis identified 19 chlorophyll a/b binding protein (CAB) genes as upregulated in the overexpressors, six of which were validated by qPCR. In wild-type cells, *CsubMADS1* was upregulated and CAB transcripts were downregulated during nitrogen starvation, indicating that both the transcription factor and these candidate downstream genes respond to nutrient stress. Six CAB genes contained CArG motifs within their 2 kb upstream regions, and CArG boxes from CAB24 and CAB8 were tested for binding. CAB24 belongs to the stress-responsive LI818 family. Agarose-based EMSA provided preliminary evidence of CsubMADS1 interaction with the CAB24 and CAB8 CArG motifs, with the enriched sequence resembling the AGL15-type C(A/T)8G motif. These observations suggest that CAB gene expression is associated with stress responses and that CsubMADS1 may regulate a subset of CABs, although additional assays are required to confirm direct regulation.

## 1. Introduction

In seed plants, extensive research on MADS-box transcription factors (TFs) has shown their role during development and stress (Kater *et al*., 2006, Castelan-Munoz *et al*., 2019). These transcription factors are gene families in the case of seed plants; for example, there are 75 in rice, 107 in Arabidopsis and 105 in Populus (Parenicová *et al*., 2003, Leseberg *et al*., 2006, Arora *et al*., 2007). Non-seed plants have MADS-box transcription factors (TF), but the numbers are much lesser than the seed plants, and most functions remain uncharacterized (Thangavel and Nayar, 2018). However, recent functional characterization of MADS-box TFs from non-seed plants has highlighted their roles during development and stress (Zobell *et al*., 2010, Koshimizu *et al*., 2018, Nayar and Thangavel, 2021). In *Marchantia*, a MIKC* gene, *MpMADS1*, binds to the DNA in a homodimeric form and is significantly redundant in function to the Arabidopsis MIKC* complexes in the pollen (Zobell *et al*., 2010). The MADS-box genes (MIKC type) from *Physcomitrium* regulate cell division in the stem required for sustaining water conduction. These genes also aid in adequately developing the motile sperm in this bryophyte (Koshimizu *et al*., 2018). A MADS-box transcription factor, CsubMADS1 from microalga *Coccomyxa* C-169 is responsible for timely autospore release and tolerance to starvation stress. It forms homodimers and consists of a functional NLS in the MADS domain. In addition to the MADS domain, CsubMADS1 harbours an I domain but lacks the K and C domains (Nayar and Thangavel, 2021). Generally, MADS-box TFs bind to the DNA as homodimers or heterodimers to a ten-base pair conserved consensus sequence, CArG box (CC(A/T)_6_GG or variant C(A/T)_8_G) (Dolan and Fields, 1991, Treisman, 1992). Little is known about the mechanism of algal MADS-box TF binding to DNA. Since there is minimal information about the functional roles of MADS-box TFs in non-seed plants, extensive research is needed to understand their evolution better.

MADS-box transcription factors are majorly homeotic genes but also function during stress (Castelan-Munoz *et al*., 2019). Several examples of MADS-box TFs participating during stress responses and tolerance exist across different plants (Gan *et al*., 2005, Gan *et al*., 2012, Khong *et al*., 2015, Yu *et al*., 2015, Chen *et al*., 2016, Guo *et al*., 2016, Chen *et al*., 2019, Nayar and Thangavel, 2021). These transcription factors are responsive to environmental cues or stress conditions and regulate the developmental processes accordingly (Castelan-Munoz *et al*., 2019). For example, in Arabidopsis, SVP in response to drought stress via ABA is responsible for inhibiting stomatal conductance and brings about changes in some developmental processes (Bechtold *et al*., 2016, Wang *et al*., 2018). In response to drought stress FLC, SVP, and SOC1 regulate flowering transition (Lee *et al*., 2007, Li *et al*., 2008, Riboni *et al*., 2013, Riboni *et al*., 2016). In Rice, *OsMADS25* is responsive to nitrate, *OsMADS26* is responsive to osmotic stress and drought, and cold stress activates *OsMADS57* to regulate growth under these stressed conditions (Khong *et al*., 2015, Yu *et al*., 2015, Chen *et al*., 2018). In microalga, *Coccomyxa*, CsubMADS1 appears to play an important role during starvation stress (Nayar and Thangavel, 2021). These studies collectively support the emerging view that MADS-box TFs play important roles during stress responses across the plant kingdom. Plant transcription factors are known to regulate cellular and developmental processes in response to nutrient-limiting conditions (Devaiah *et al*., 2007, Yang *et al*., 2016, Baek *et al*., 2017, Guan *et al*., 2017, Luang *et al*., 2018, Barragán-Rosillo *et al*., 2021, Ried *et al*., 2021, Das *et al*., 2022, Shibata *et al*., 2022). In microalgae also, transcription factors control different cellular processes in nutrient-limiting conditions. For example, PSR1 (Phosphorus starvation Response 1), an MYB-type transcription factor upregulated during phosphorus starvation, is present in *Chlamydomonas, Nannochloropsis* and *Phaeodactylum.* PSR1 is responsible for adapting the microalgae to an environment lacking phosphorus by affecting membrane lipid profiles, starch metabolism, P scavenging, phospholipid remodeling, and cell growth (Wykoff *et al*., 1999, Bajhaiya *et al*., 2016, Kumar Sharma *et al*., 2020, Murakami *et al*., 2020). Lipid Remodelling Regulator 1 (LRL1) from *Chlamydomonas* plays an essential role during P-deprivation by regulating genes required for the cell cycle, phosphorus, and lipid metabolism (Hidayati *et al*., 2019). Transcription factors like nsbHLH2 from *Nanochloropsis*, ROC40 from *Chlamydomonas*, CmMYB1 from *Cyanidioschyzon* and bZIP14 from *Phaeodactylum* all play pivotal roles in the nitrogen-depleted conditions (Imamura *et al*., 2009, Kang *et al*., 2015, Goncalves *et al*., 2016, Matthijs *et al*., 2017). Copper response regulator 1 (CRR1) from *Chlamydomonas* is a global regulator of genes involved in copper deficiency and has a role in zinc homeostasis (Lambertz *et al*., 2010, Sommer *et al*., 2010). Another TF from Chlamydomonas, FEMU2, regulates FOX2, which is responsible for cellular Fe uptake during iron deficiency (Deng *et al*., 2014). These reports reaffirm the role of transcription factors in macronutrient and micronutrient-limiting conditions.

Our previous study showed that CsubMADS1, a MADS-box TF from *Coccomyxa* C-169, has a role in starvation stress (Nayar and Thangavel, 2021). In this study, further work has been done to show its function during nutrient-limiting conditions, such as stationary phase and nitrogen deprivation, using the *CsubMADS1* overexpressors. Possible direct downstream targets of CsubMADS1 have been identified using RNA Seq and qPCR. Further, electrophoretic mobility shift assays were performed to examine the potential binding of CsubMADS1 to the regulatory element in the target genes.

## 2. Results and Discussion

### 2.1 *CsubMADS1* overexpression is associated with increased tolerance to nitrogen starvation

Previous work from our lab showed that overexpressors of *CsubMADS1* in *Coccomyxa* C-169 had reduced mucilage production during starvation (Nayar and Thangavel, 2021). Therefore, the present study revolves around probing for the role of CsubMADS1 in stress tolerance during starvation. *CsubMADS1* overexpressors (OX1, OX5 and OX7) delineated in the previous study were subjected to nitrogen starvation. These lines were compared to the wild type grown under nitrogen starvation. MADS OX1 and OX5 were tolerant to nitrogen starvation compared to the wild type, whereas OX7 was similar to the wild type (Figure 1A, Figure S1). Thus, OX1 and OX5 lines have been used for comparison to the wild type for further experiments. In addition, the status of the *CsubMADS1* transcript was checked under nitrogen-sufficient and nitrogen-deficient conditions in the wild type. *CsubMADS1* transcript was upregulated almost two-fold in the nitrogen-deficient state (Figure 1B). In OX1 and OX5, *CsubMADS1* was upregulated in nitrogen-sufficient and nitrogen-deficient conditions (Figure 1C). Thus, these results suggest that CsubMADS1 is associated with increased stress tolerance and is responsive to nitrogen starvation.

**Figure 1.**
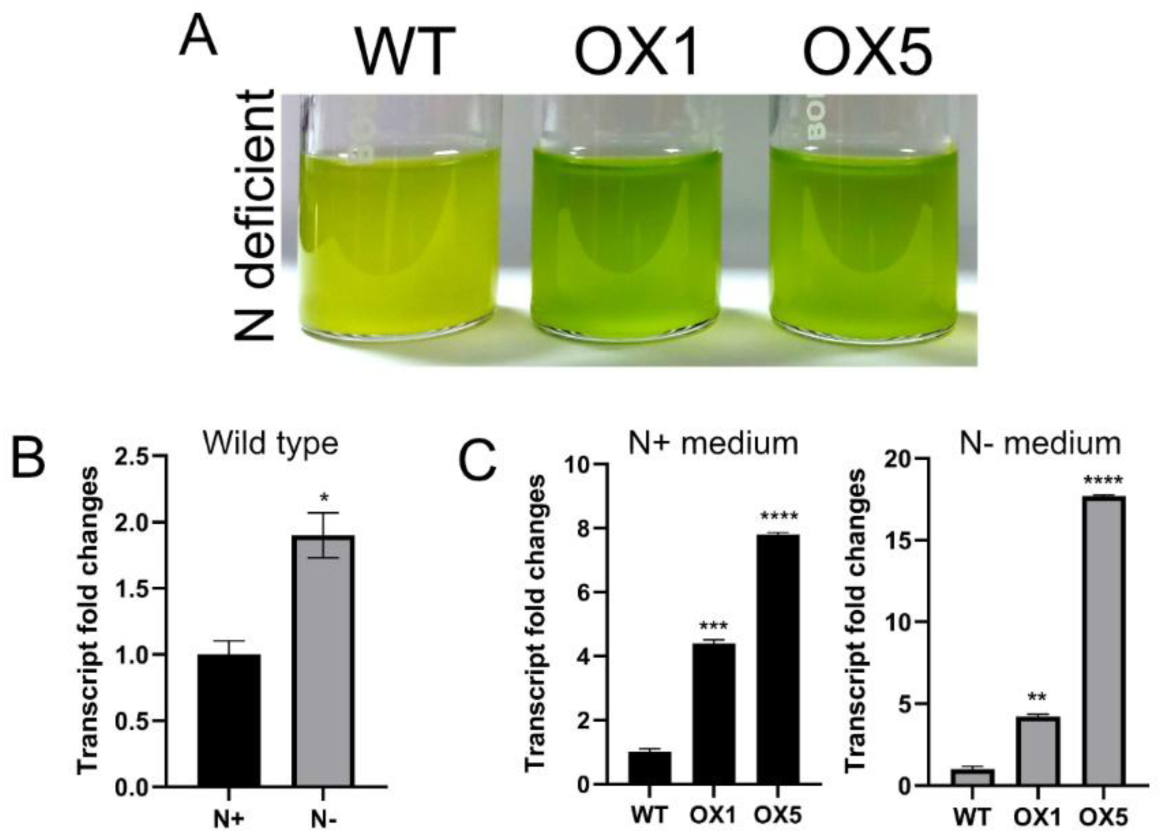
CsubMADS1 overexpressors are tolerant to nitrogen starvation A) Comparison of wild-type C-169 and CsubMADS1 OX1 and OX5 cultures grown in nitrogen-deficient medium. The overexpressors show better tolerance to the nitrogen deficiency than the wild-type B) Quantitative PCR; transcript abundance of CsubMADS1 in wild-type C-169 in N-sufficient versus deficient medium. Bar indicates ±SE (n = 3). C) Quantitative PCR; Left panel; Transcript abundance of CsubMADS1 in wild-type versus CsubMADS1 overexpressors (OX1 and OX5) grown in N-sufficient medium. Right panel; Transcript abundance of CsubMADS1 in wild-type versus CsubMADS1 overexpressors (OX1 and OX5). Bar indicates ±SE (n = 3). For all graphs, data were significant at ****P ≤ 0.0001, ***P ≤ 0.001, **P ≤ 0.01, and *P ≤ 0.05 compared with the wild type using Student’s t-test.

MADS-box transcription factors, *ANR1* from Arabidopsis and *OsMADS61* from Rice, belong to the *AGL17*-like clade and are induced by nitrogen starvation (Gan *et al*., 2005, Yu *et al*., 2014). In addition, in Arabidopsis, other genes from this clade (*AGL14, AGL16, AGL19, SOC1, AGL21, AGL26* and *AGL56*) are also induced by nitrogen starvation (Gan *et al*., 2005). In rice, *OsMADS25* and *OsMADS27* were induced by nitrate resupply, whereas *OsMADS57* was downregulated by nitrogen deprivation (Yu *et al*., 2014). The genes in the *AGL17*-like clade are involved in root development, and most are responsive to changes in nitrogen, phosphorus, and sulphur levels (Gan *et al*., 2005, Gan *et al*., 2012, Yu *et al*., 2014). *CsubMADS1*, like *OsMADS61* and *ANR1*, is upregulated during nitrogen starvation. But it remains to be studied whether the *CsubMADS1* transcript is altered due to fluctuation of other nutrients, for example, phosphorus and sulphur. Previous research on *Chlamydomonas* showed that the alga undergoes nitrogen deprivation during cold stress (Valledor *et al*., 2013). *Coccomyxa* C-169 is a cold-adapted microalga first isolated from Antarctica (Blanc *et al*., 2012). Hence a pertinent question regarding the role of CsubMADS1 is whether it has any role during cold adaptation.

### 2.2 Stress tolerance of CsubMADS1 OX lines

To further investigate the role of CsubMADS1 during stress tolerance, the wild-type and overexpressors were grown in a nitrogen-sufficient medium until the stationary phase. The status of the neutral lipids was checked during these conditions. The wild-type and overexpressors were stained with Nile red, and the abundance of lipid droplets was determined by fluorescence-assisted cell sorting (FACS) and confocal imaging. In the wild type, in a nitrogen-sufficient medium, Nile red-specific fluorescence was detected in almost 32% of the cell population (Figure 2A, B). The confocal imaging also corroborated the high number of lipid droplets in the wild type (Figure 2A). However, in the overexpressors in the same medium, fluorescence was detected only in two per cent of the cell population (Figure 2A, B). The same result was also evident in confocal microscopy (Figure 2A). The cells were also stained with methylene to detect mucilage production. The wild-type cells were stained blue, and overexpressor cells were visible as a mix of green and patches of blue (Figure 2C). Decreased staining of overexpressors with methylene revealed that mucilage production in the overexpressors is lower than in the wild type. These observations are consistent with a reduced stress-associated lipid accumulation in the overexpressors, although additional assays would be required to establish a connection. In *Phaeodactylum tricornutum*, similar results were seen where an increase in lipid droplets was observed during the stationary phase in replete conditions (Cavonius *et al*., 2015).

**Figure 2.**
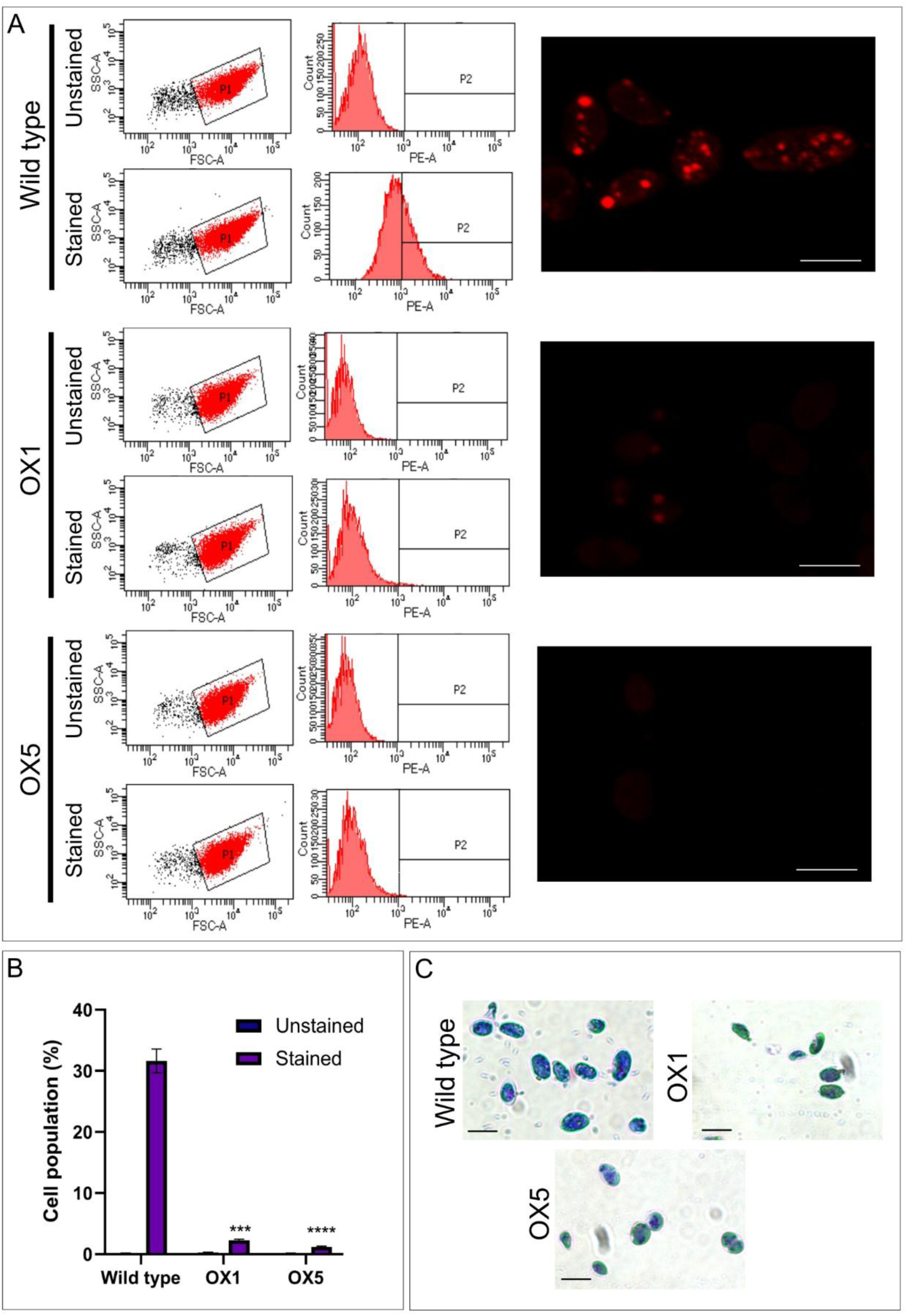
Decreased stress response in overexpressors during stationary phase. A) Wild type and overexpressors (OX1 and OX5) at their stationary phase were stained with Nile red and subjected to flow cytometry to detect neutral lipids in their cell populations. Unstained cells were used as a control. The stained cells were also observed under the confocal microscope to detect the lipid droplets. B) Neutral lipids detected using flow cytometry in the Nile red stained cells of Wild type, OX1 and OX5 from the stationary phase were plotted in terms of percentage. ±SE (n = 3).C) Methylene staining of cells of WT, OX1, and OX5 at stationary phase to detect mucilage on the cell surface; Bar= 5 μm. For all graphs, data were significant at ****P ≤ 0.0001 and ***P ≤ 0.001 when compared with the wild type using Student’s t-test.

Additionally, wild-type and overexpressors were grown in a nitrogen-deficient medium until the stationary phase. Finally, the lipid droplets stained with Nile red were quantified using FACS and visualized using confocal microscopy. It is known from several previous studies that nitrogen starvation induces increased lipid droplet formation in microalgae (Msanne *et al*., 2012, Fan *et al*., 2014, Cavonius *et al*., 2015, Goncalves *et al*., 2016). In the wild-type, Nile red was detected in 53% of the cell population (Figure 3A, B), which was also visible in the confocal microscopy. However, in overexpressors, Nile red was detected only in two per cent of the cell population even after nitrogen starvation (Figure 3 A, B), which was confirmed by confocal microscopy. Furthermore, wild-type and overexpressor were stained with methylene blue to check mucilage formation. Even in the nitrogen-deficient medium, methylene staining was decreased in the overexpressors than in the wild-type (Figure 3C). These observations indicate reduced mucilage staining in the overexpressors under the tested conditions. A 21% increase of lipid droplets was seen in stationary phase wild-type cells grown in a nitrogen-deficient medium versus a nitrogen-sufficient medium. In contrast, the percentage of lipid droplets remained the same in the overexpressor in both media. Hence, *CsubMADS1* overexpression phenotype in both nitrogen-sufficient and nitrogen-deficient media at the stationary phase showed altered physiological responses typically associated with stress however, additional assays are needed to directly link these traits to stress tolerance.

**Figure 3.**
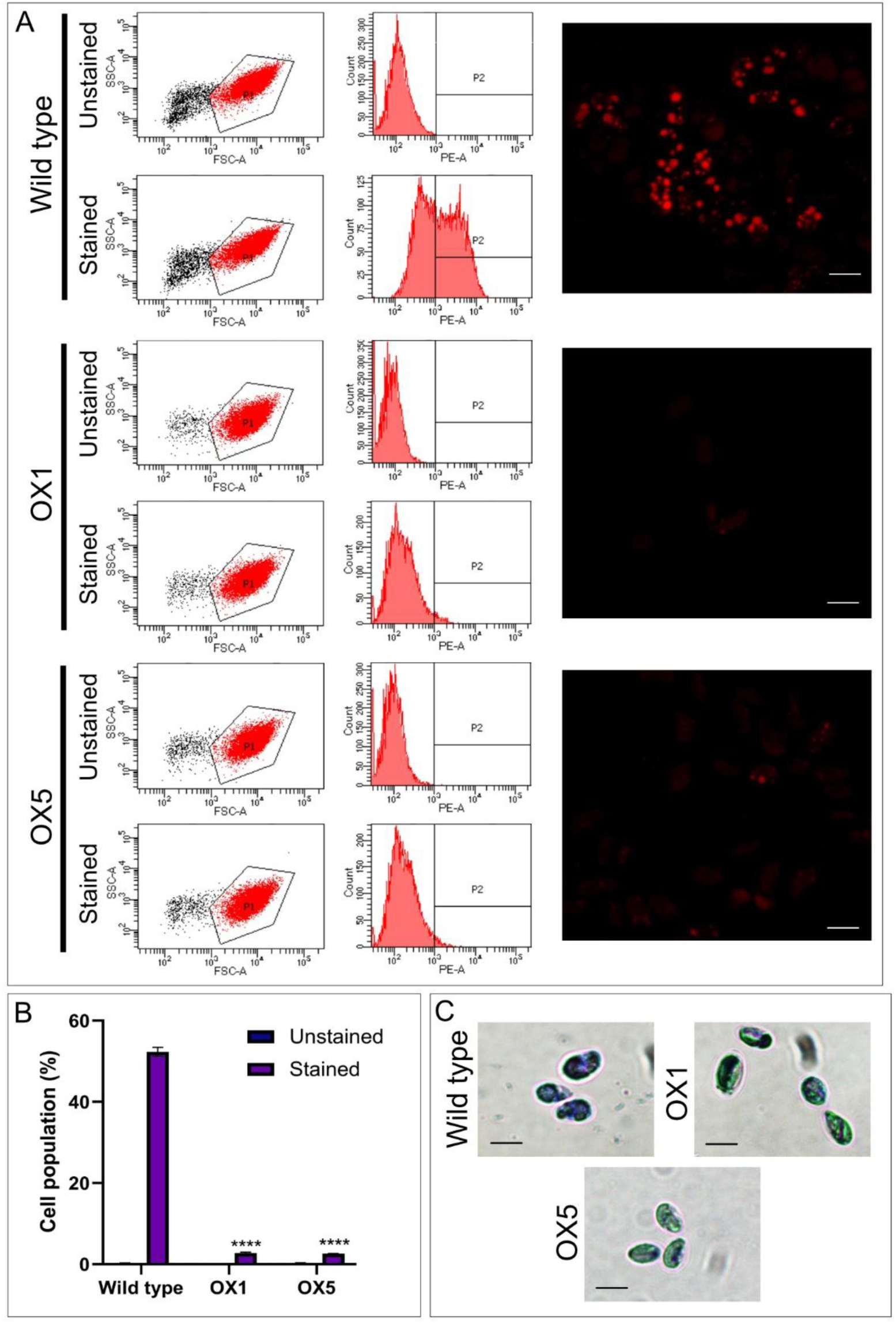
Reduced stress response in overexpressors during nitrogen starvation. A) Cells from nitrogen-starved cultures of wild-type and overexpressors (OX1 and OX5) were stained with Nile red and subjected to flow cytometry to detect neutral lipids in their cell populations. Unstained cells were used as a control. The stained cells were also observed under the confocal microscope to detect the lipid droplets. B) Neutral lipids detected using flow cytometry in the Nile red stained cells of wild type, OX1 and OX5 from nitrogen starved cultures were plotted in terms of percentage. ±SE (n = 3). C) Methylene staining of cells from nitrogen-starved cultures of WT, OX1, and OX5 to detect mucilage on the cell surface; Bar= 5 μm. For all graphs, data were significant at ****P ≤ 0.0001 when compared with the wild type using Student’s t-test.

Several MADS-box transcription factors are stress-responsive and are a crucial component of stress response in seed plants to abiotic stresses (Castelan-Munoz *et al*., 2019). For example, *OsMADS25* is induced in rice by high nitrate levels that control primary and lateral root growth (Yu *et al*., 2015). *OsMADS26* is responsive to osmotic and drought stress to regulate tiller growth (Khong *et al*., 2015). Additionally, *OsMADS82* and *OsMADS87* are inhibited by heat stress and are involved during seed development (Chen *et al*., 2016). A gene from pepper, *CaMADS*, is responsive to cold, osmotic and salt stress (Chen *et al*., 2019). SlMBP11 from tomato is a drought, wounding, and salt stress-responsive gene responsible for seedling growth (Guo *et al*., 2016). Our previous study showed that CsubMADS1 may contribute to starvation stress responses in *Coccomyxa* C-169 (Nayar and Thangavel, 2021). In this study, we have used stationary phase and nitrogen deprivation as nutrient-limiting conditions to elucidate its role during starvation stress. The examples mentioned above show that a single MADS-box gene can be responsive to multiple abiotic stresses; thus, it remains to be studied whether CsubMADS1 is partaking in regulating other abiotic stress response pathways or not. CsubMADS1 is a lag phase transcription factor, the phase during which microorganisms are exposed to cellular stress (Hamill *et al*., 2020). Hence, the proposed role of CsubMADS1 in stress responses is consistent with these observations, although further investigation is needed to clarify its specific function.

### 2.3 Chlorophyll A/B binding protein (CAB) genes are a plausible downstream target of CsubMADS1

To find the possible downstream targets of CsubMADS1, transcriptomic analysis was done using RNA seq from the stationary phase of wild-type and overexpressors grown in nitrogen sufficient medium, with three biological replicates each. In the RNA Seq, the percentage of reads aligned to the reference genome was around 65.79% which is typical for non-model microalgae given incomplete genome annotations, strain-level variation, and the presence of organellar and non-coding RNA species. In many experiments, alignment rates of 70–95% are common, although the exact values depend strongly on sample quality, RNA integrity, genome completeness, and alignment parameters. Importantly, in this experiment, each sample retained >10 million uniquely aligned reads, providing adequate depth for differential expression analysis. There were 657 genes downregulated and 818 genes upregulated with 1.5-fold change and p-value ≤0.05 according to the differential expression analysis in the overexpressor vs. wild-type (Figure 4A). According to RNA Seq data, *CsubMADS1* is upregulated in the overexpressor by 1.8-fold. The RNA Seq data was also validated by qPCR. According to qPCR, *CsubMADS1* is upregulated by four and eightfold in OX1 and OX5, respectively.

**Figure 4.**
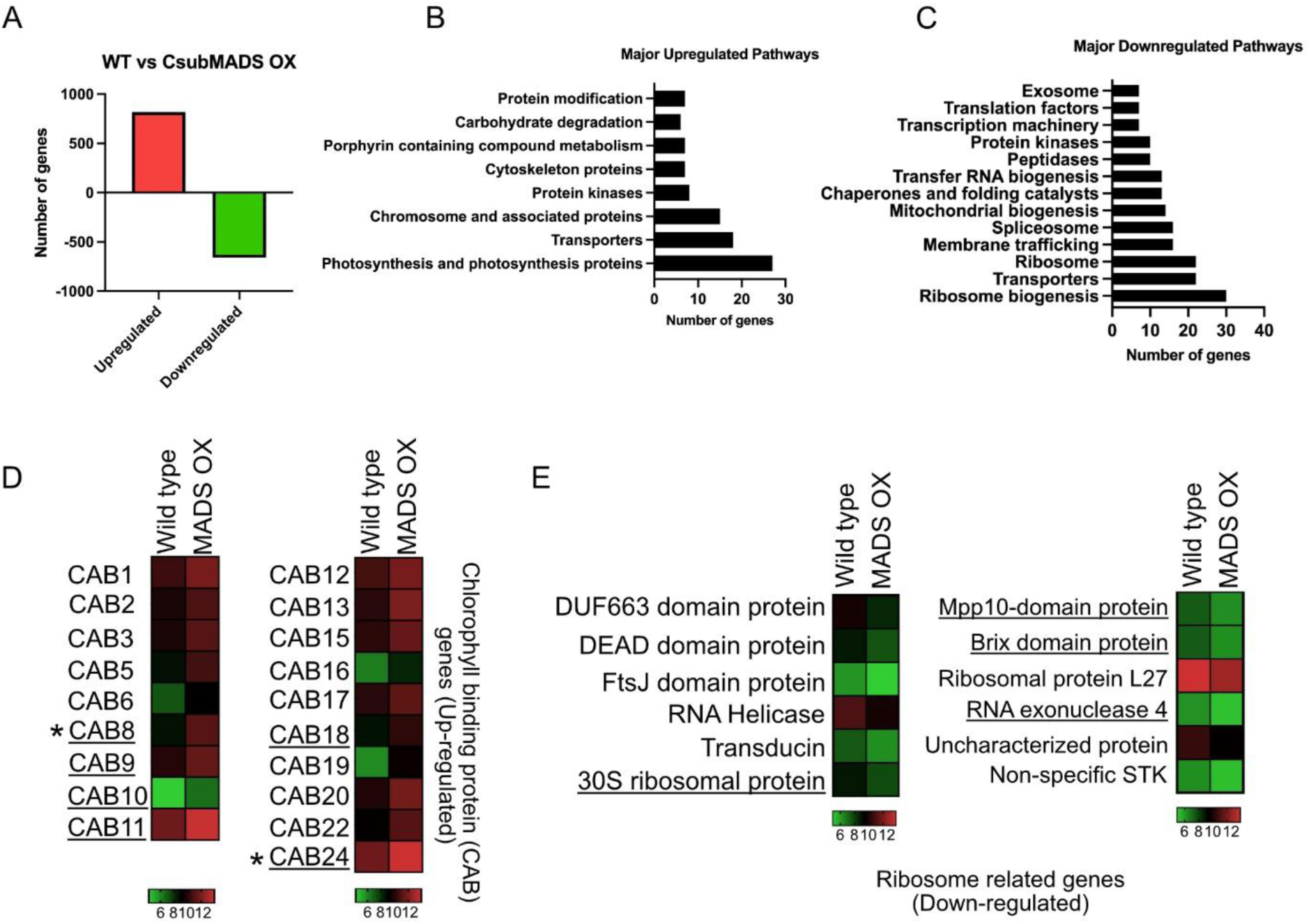
RNA-Seq based comparison of *CsubMADS1* OX and WT transcriptome at stationary phase (A) Total number of differentially expressed genes (1.5 Fold) in *CsubMADS1* OX compared to the wild type; upregulated genes (red) and downregulated genes (green) (pValue ≤ 0.05, n = 3). (B) Major upregulated pathways in *CsubMADS1* OX in comparison to WT. (C) Major downregulated pathways in *CsubMADS1* OX compared to WT. (D) Heat maps of a selected upregulated pathway (photosynthesis-related genes) in *CsubMADS1* OX; colour bars below each sub-panel represent the range of expression values in log_2_. (E) Heat maps of a selected downregulated pathway (ribosome-related genes) in *CsubMADS1* OX; colour bars below each sub-panel represent the range of expression values in log_2_. Underlined genes in the heatmap have a CArG box in their 2 kb upstream region. The CAB genes marked with * were selected for further studies.

Further, the major upregulated and downregulated pathways were identified. In the upregulated pathways, photosynthesis-related genes were the top category, whereas, in the downregulated pathways, ribosome-related genes were the highest in number (Figure 4 B, C, Table S1, Table S2). 19 chlorophyll A/B binding protein (*CAB*) genes were upregulated, belonging to the photosynthesis-related genes (Figure 4D, Table S3). Moreover, six *CAB*s had a CArG motif in their 2 kb upstream region (Figure S2), suggesting they maybe potential candidates for regulation by CsubMADS1. Some downregulated ribosomal-related genes, such as MPP10-domain protein, RNA exonuclease 4, Brix domain protein, and 30S ribosomal protein genes, also had CArG motifs in their 2 kb upstream region (Figure 4E) indicating that they too may be candidates for transcriptional regulation.

In algae and higher-order plants, chlorophyll-binding proteins respond primarily to light but additionally, they are also responsive to various other signals and stresses (Silverthorne and Tobin, 1984, Millar and Kay, 1996, Andersson *et al*., 2003, Varsano *et al*., 2006, Hwang *et al*., 2008, Staneloni *et al*., 2008, Dittami *et al*., 2009, Dittami *et al*., 2010, Park *et al*., 2010). In Arabidopsis, *LHCB/CAB* genes are responsive to abscisic acid (ABA), and the downregulation of at least six of these makes the plant insensitive to ABA. A transcription factor, WRKY40, a repressor, is also involved in balancing *LHCB/CAB* functions (Liu *et al*., 2013). In another study on Arabidopsis *CAB* genes, the downregulation of these genes caused decreased sensitivity of stomata to ABA, resulting in reduced drought stress tolerance in the plants (Xu *et al*., 2012). In *Camellia sinensis*, a CAB gene, *CsCP1*, is downregulated in response to six stress conditions (Li *et al*., 2020). In jatropha, CAB genes *LHCB*s are responsive to hormones, drought and salt stress (Zhao *et al*., 2020). Small CAB-like proteins (*Scp*) from *Synechocystis*, *ScpB-ScpE* expression upregulated in response to various stress conditions and a mutant lacking four Scp’s lacked tolerance to high light conditions (He *et al*., 2001). Chlorophyll a/c binding proteins (*FCP*) from *Ectocarpus* are also responsive to various stresses (Dittami *et al*., 2009). In the *CsubMADS1* overexpressor, 19 *CAB* genes were upregulated, and the overexpressor exhibited tolerance to starvation, raising the possibility that some CABs could contribute to this phenotype, although direct functional connections remain to be established.

Ribosomal proteins participate in translation and have alternative functions during stress responses (Wool, 1996, Warner and McIntosh, 2009). For example, overexpression of *RPL6* in rice conferred salt stress tolerance (Moin *et al*., 2020), and the knockdown of a 60S ribosomal protein *L14-2* in cotton boosted drought and salt tolerance in these plants (Shiraku *et al*., 2021). The silencing of *RPL12* and *RPL19* in *N. benthamiana* resulted in a lag of the nonhost bacteria-induced hypersensitive response (HR) and increased growth of nonhost bacteria, indicating that ribosomal proteins also play a role during biotic stress (Nagaraj *et al*., 2016). Ribosome biogenesis decreases when *RUNX1* is mutated, giving stress resistance to hematopoietic stem and progenitor cells (Cai *et al*., 2015). In *CsubMADS1* overexpressors, the ribosome biogenesis-related genes were downregulated, which may reflect a shift in metabolic prioritization during nutrient limiting conditions, but the mechanism remains uncertain. Since there is a known role for *CABs* during stress tolerance and 19 were upregulated in the overexpressors, their possible involvement in starvation tolerance was explored further.

### 2.4 Chlorophyll A/B binding protein (CAB) gene expression is altered during nitrogen starvation

*Coccomyxa* C-169 has 24 Chlorophyll A/B binding protein (*CAB*) genes, of which *CAB24* belongs to the LI818 family (Dittami *et al*., 2010), a family reported to be involved in stress responses (Table S4). Out of 19 upregulated CAB genes, the abundance of six *CAB* transcripts, three genes with CArG boxes in the upstream region; *CAB24, CAB8, CAB9* and three genes without CArG boxes in the upstream region; *CAB20, CAB15*, and *CAB17*, was studied using qPCR in wild-type in a nitrogen-sufficient and deficient medium. These analyses indicated *CAB*s were downregulated in response to nitrogen starvation in the wild-type (Figure 5). In addition, the abundance of the chosen *CAB* genes was also studied in overexpressors (OX1, OX5) and compared to wild-type in both nitrogen-sufficient and nitrogen-deficient media. *CAB*s were upregulated in the overexpressors compared to the wild type in nitrogen-sufficient and nitrogen-deficient media (Figure 6). In the nitrogen-sufficient medium, *CAB*s were upregulated by approximately 3 to 10-fold in the overexpressor compared to the wild type. In the nitrogen-deficient medium, *CAB*s were upregulated around 5 to 50-fold in the overexpressor compared to the wild type. The larger fold-change observed under nitrogen deficiency likely reflects the reduced baseline *CAB* transcript levels in the wild type. *CAB24* and *CAB8* were highly upregulated in the nitrogen-deficient condition and possessed CArG boxes in their upstream region; hence these genes were selected for further experiments.

**Figure 5.**
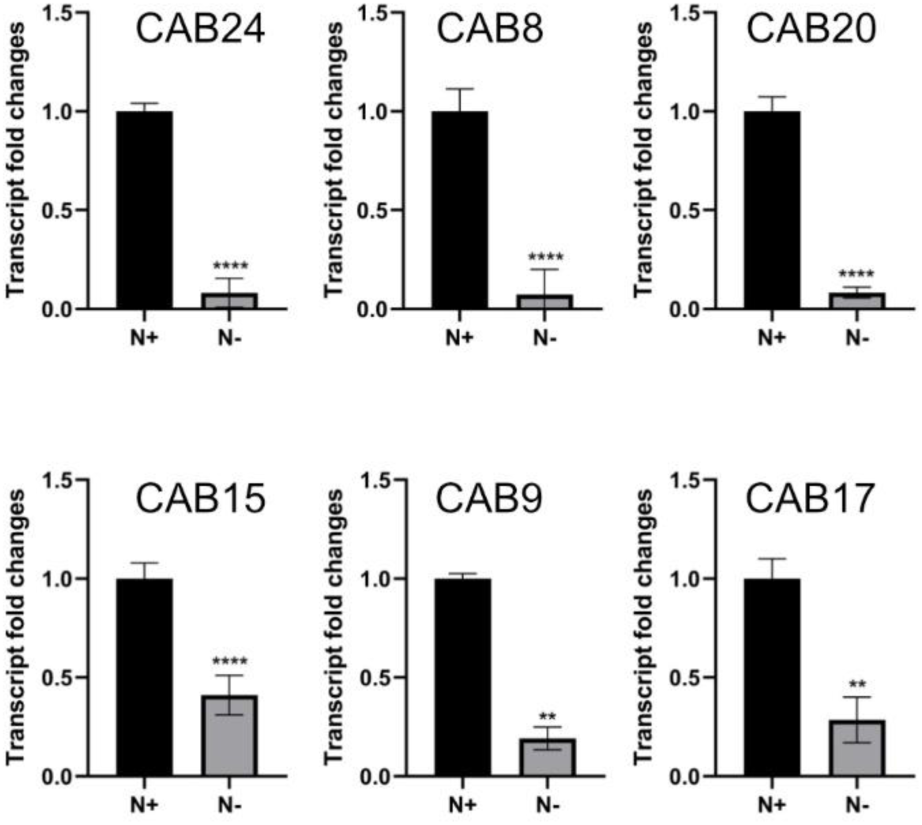
Chlorophyll A/B binding proteins (*CsubCAB*) genes downregulated during nitrogen starvation. Quantitative PCR; This panel shows the transcript abundance of six *CsubCAB*s in wild-type grown in N-sufficient versus N-deficient medium. Bar indicates ±SE (n = 3). For all graphs, data were significant at ****P ≤ 0.0001 and **P ≤ 0.01 compared with the wild-type N-sufficient using Student’s t-test.

**Figure 6.**
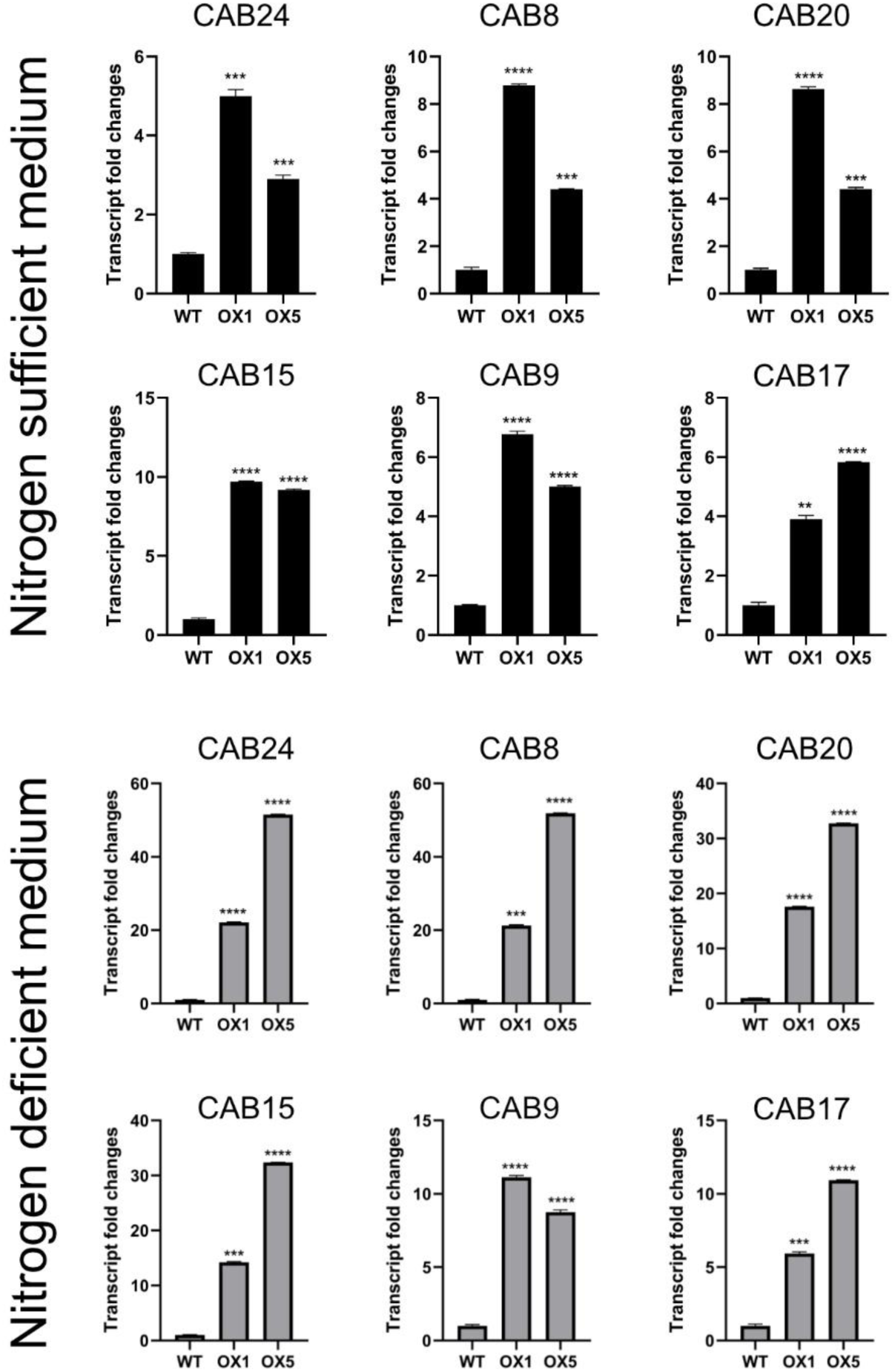
Chlorophyll A/B binding proteins (*CsubCAB*) genes are upregulated in the overexpressors. Quantitative PCR; Upper panel shows the transcript abundance of six *CsubCAB*s in wild type and overexpressors grown in an N-sufficient medium. The lower panel shows the transcript abundance of six *CsubCAB*s in the wild-type and overexpressors grown in the N-deficient medium. Bar indicates ±SE (n = 3). For all graphs, data were significant at ****P ≤ 0.0001, ***P ≤ 0.001 and **P ≤ 0.01 compared with the wild type using Student’s t-test.

Nitrogen starvation effectively directs the metabolites to oil production in microalgae (Msanne *et al*., 2012, Fan *et al*., 2014, Goncalves *et al*., 2016). Though nitrogen starvation enhances lipid production, it has a detrimental effect on microalgae growth as it damages the photosynthetic apparatus and diminishes the light-harvesting ability of the algae (Krapp *et al*., 2011, Møller *et al*., 2011, Wu *et al*., 2022). This inability to fix carbon efficiently has been proposed to contribute to activating the lipid pathways, thereby upregulating lipid metabolism genes (Goncalves *et al*., 2016). Chlorophyll A/B binding proteins are accessory proteins of the light-harvesting complex, which have also been implicated in stress responses (Dittami *et al*., 2010) but their role during nitrogen starvation is still not fully understood. For example, a study in the diatom, *Phaeodactylum tricornutum,* showed a chlorophyll-binding protein, fucoxanthin-chlorophyll a/c binding protein B (*FCP B*), is severely downregulated during nitrogen starvation conditions (Curcuraci *et al*., 2022). A similar trend was observed in the present study in wild-type. Also, another study in Arabidopsis reported that a mutant overexpressing a chlorophyll A/B binding protein (*COE2)* showed increased tolerance to nitrogen starvation compared to the wild type (Wu *et al*., 2022). Similarly, in *CsubMADS1* overexpressors, *CAB* transcript upregulation correlated with the observed tolerance to nitrogen starvation. Hence, these findings suggest potentially similar transcriptional responses of *CAB* in *P. tricornutum* (diatom), Arabidopsis (dicot) and *C. subellipsoidea* (green microalgae) during nitrogen starvation, though the underlying mechanisms remain distinct and require further study.

### 2.5 CsubMADS1 shows indicative binding to CArG motif–containing probes under the agarose-based EMSA conditions

Since many *CAB* genes were upregulated in the *CsubMADS1* overexpressors, the 2 kb upstream regions of the 19 upregulated *CAB* genes were scanned for CArG motifs using the PLACE database. This analysis showed six *CAB* genes had CArG motifs in their upstream region, which shows that probably these genes may be the potential targets of this transcription factor (Table S5). This study did not consider the CArG boxes in other potential regulatory regions of the gene, like 5’UTR and introns. CArG motifs of the *CAB24* and *CAB8* upstream regions were chosen for EMSA studies to determine whether CsubMADS1 could bind to these motifs. Recombinant His-tagged CsubMADS1 was purified, and the presence of the protein was confirmed in the whole lysate and purified fractions by western analysis using anti-His and anti-CsubMADS1 antibodies (Figure 7A, Figure S3). *CAB24* and *CAB8* CArG DNA probes were tested for binding with CsubMADS1 protein in a dose-dependent manner (500 ng to 2000 ng). Recombinant CsubMADS1 exhibited binding to *CAB24* and *CAB8* CArG motif probes in EMSA assays, with a noticeable mobility retardation upon addition of protein. (Figure 7B, C). Increasing CsubMADS1 protein amount resulted in progressively slower-migrating complexes, suggesting formation of possibly larger protein–DNA assemblies, although the exact nature of these complexes cannot be determined solely from migration patterns. Progressively slower migration of DNA-protein complexes with increasing protein concentration has been observed in both PAGE- and agarose-based EMSAs, although the exact banding patterns differ depending on gel matrix and complex size (Jing *et al*., 2003, Cordeiro *et al*., 2011, Wilkinson *et al*., 2019).

**Figure 7.**
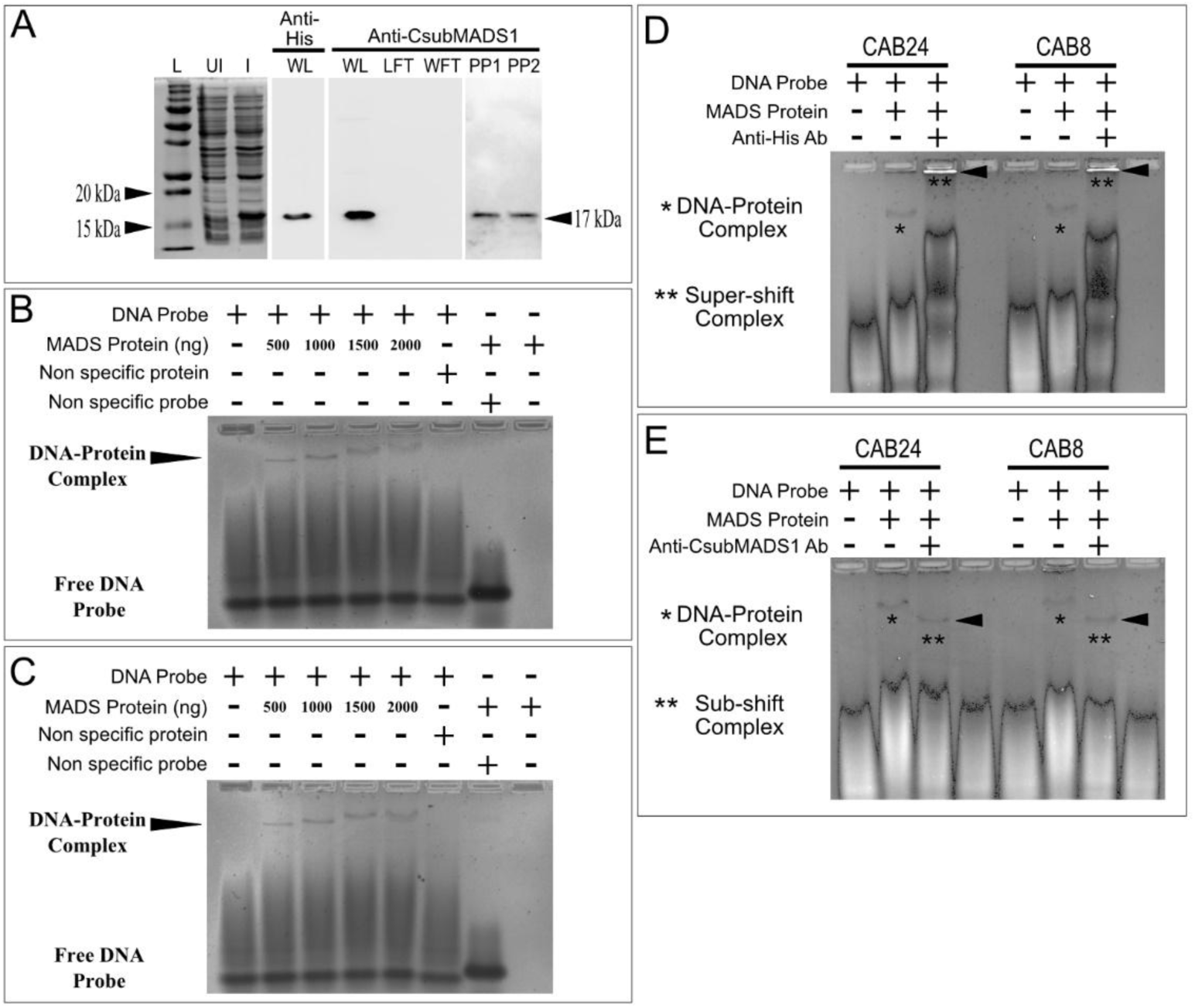
EMSA suggests that CsubMADS1 may interact with CArG motifs in CAB24 and CAB8; however, these interactions should be interpreted cautiously due to the limitations of agarose-based EMSA. A) First panel; SDS-PAGE shows uninduced (UI) and induced (I) lysate of E. coli expressing CsubMADS1 protein and protein ladder. Second panel; Western blot shows CsubMADS1 band probed with anti-His antibody in the whole lysate. Third panel; Western blot shows CsubMADS1 band probed with anti-CsubMADS1 antibody in the whole lysate. No band was seen in the lysate flow through (Krapp *et al*.) and wash flow through (WFT). Fourth panel; Western blot shows CsubMADS1 band probed with anti-CsubMADS1 antibody in the purified protein (PP) fraction. B) Electromobility shift assay (EMSA) agarose gel shows binding of CsubMADS1 to the CArG box of *CAB24*. An increasing amount of the protein, 500 ng to 2000 ng, caused increased gel retardation. C) EMSA agarose gel displays binding of CsubMADS1 to the CArG box of *CAB8*. An increasing amount of the protein, 500n g to 2000 ng, caused increased gel retardation. D) Super shift EMSA agarose gel reveals binding of the anti-His antibody to the CsubMADS1-CArG complex for *CAB24* and *CAB8*. E) EMSA gel displays sub shift of DNA-protein complex for *CAB24* and *CAB8*, which may be due to the sequestration of CsubMADS1 by the anti-CsubMADS1 antibody. A peptide from the DNA binding domain was used to raise the antibody. The antibody binding to the CsubMADS1 DNA binding domain may prevent binding to the CArG box.

To further validate these results, super shift EMSAs were performed using anti-His and anti-CsubMADS1 antibodies. CsubMADS1 bound to *CAB24* and *CAB8* probe saw a supershift in the presence of anti-His antibody (Figure 7D). But in contrast, the anti-CsubMADS1 antibody caused the DNA-protein complex to sub-shift (Figure 7E). The subshift likely occurred because the epitope (RIEKIGDERNRQVTFTKRKN) recognized by the CsubMADS1 antibody was present in the DNA binding domain of the CsubMADS1 protein. This was confirmed by sequence search of the DNA binding domain of CsubMADS1 (Table S6). This may have led to the CsubMADS1 antibody sequestering CsubMADS1 and occupying the DNA binding region, thus preventing it from binding to the DNA probe but additional assays would be needed to confirm this interpretation. A previous study has shown this type of subshift in EMSA experiments and attributed it to the inability of the antibody-protein complex to attach to the DNA probe (Hanakahi *et al*., 1997). Thus, supershift and subshift assays using anti-His and anti-CsubMADS1 antibodies altered the migration of the DNA–protein complex, supporting the involvement of CsubMADS1 in the observed shift. However, given the use of agarose gels and absence of competition assays, these EMSA results should be interpreted as indicative rather than definitive evidence of direct binding.

MADS-box transcription factors bind to conserved CArG motifs on the DNA, especially the CC(A/T)_6_GG type (West *et al*., 1997). The C(A/T)_8_G type is another form in seed plants with a more extended A/T-rich core sequence first shown to be bound by AGL15 (Tang and Perry, 2003). The CArG motifs enriched for CsubMADS1 binding to the upstream region of *CAB*s were the AGL15 type CArG motif. Therefore, CsubMADS1, is likely capable of interacting with CArG motifs, consistent with other MADS-domain proteins. Our previous study showed that CsubMADS1 forms homodimers in-vivo (Nayar and Thangavel, 2021) and may interact with DNA in this form. It is also interesting to note that CAB24 belongs to the LI818 family of chlorophyll-binding proteins, which have be associated with stress responses (Dittami *et al*., 2010). Six of the 19 *CAB* genes upregulated in the overexpressor had CArG boxes in their upstream region. The other *CAB* genes may be regulated indirectly, or the CArG boxes may be present in other regulatory regions. Thus, CsubMADS1 may regulate *CAB24* and *CAB8* through interaction with the CArG boxes although further validation such as competition assays, PAGE-based EMSA, and chromatin immunoprecipitation (ChIP-qPCR) will be required to confirm direct regulation. Thus, upregulation of the *CAB* genes in the overexpressors appears to be associated with stress tolerance during nutrient-limiting conditions such as stationary phase and nitrogen starvation.

## 3. Conclusions

In summary, the results presented here give insight into the functional aspect of CsubMADS1 during starvation stress (Figure 8). The overexpressors appeared to withstand the stressed condition better than the wild type. *CsubMADS1* is upregulated during nitrogen starvation, and *CAB*s are downregulated during nitrogen starvation. Furthermore, *CAB* upregulation may contribute to the physiological differences observed, although the relationship remains correlative. CsubMADS1 showed in-vitro binding to the C(A/T)_8_G type CArG motifs of *CAB24* and *CAB8* under the assay conditions. These results suggest a potential direct interaction but will require PAGE-EMSA and ChIP for confirmation. The regulation of other *CAB* genes may be indirect, or the CArG motif may be present in other regulatory regions, which also warrants further exploration. Previous studies have suggested that lag phase can involve physiological adjustments, including responses to cellular stress in some microbes (Hamill *et al*., 2020). Since *CsubMADS1* is expressed strongly during the lag phase this is consistent with, though does not by itself demonstrate, a role in stress-associated transcriptional changes.

**Figure 8.**
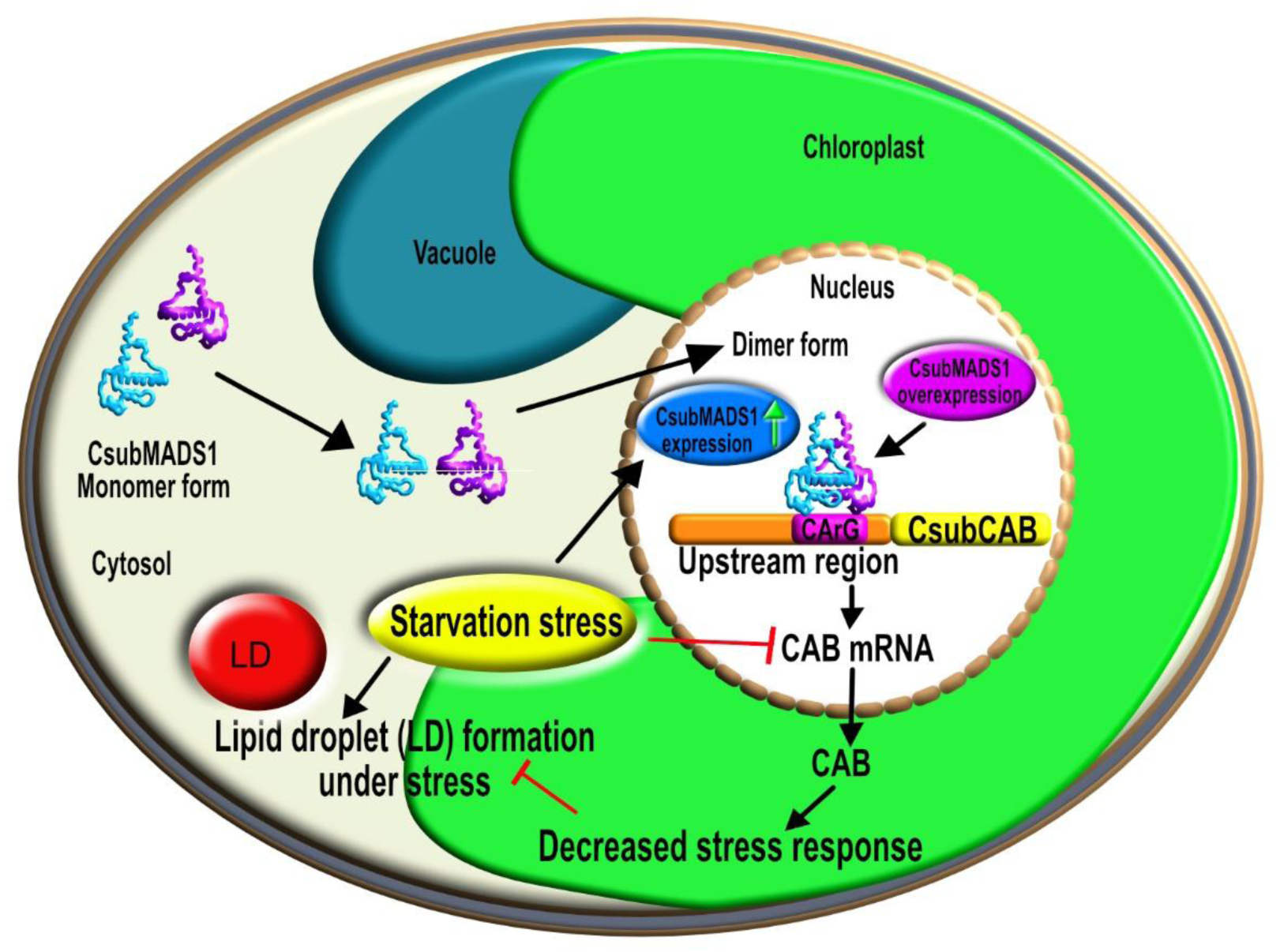
Model for CsubMADS1 role during starvation stress based on correlative evidence and proposed interactions. Our previous study showed that CsubMADS1 localizes to the nucleus as a homodimer. We also know mucilage production during starvation stress was negatively regulated in the lines overexpressing CsubMADS1. In this study, the role of CsubMADS1 during starvation stress was examined further using overexpressors at the stationary phase and treated with nitrogen deprivation. Starvation or nutrient stress induces lipid droplet formation, which was highly reduced in the overexpressors; hence CsubMADS1 negatively regulates starvation-induced lipid droplet formation. *CsubMADS1* gets upregulated during nitrogen deprivation, whereas *CAB* genes are downregulated in response to nitrogen deprivation. This transcription factor can bind to the CArG boxes in the upstream region of two *CAB* genes. In the overexpressors, high expression of CsubMADS1 led to upregulation of *CAB* genes. *CAB* upregulation was associated with decreased lipid droplet accumulation, a marker of reduced stress response. Therefore, CsubMADS1 appears to contribute to stress responses in *Coccomyxa*.

However, questions remain about whether *CsubMADS1* is responsive to other abiotic stresses, especially cold stress, as *Coccomyxa* C-169 is a microalga that can survive Antarctica’s cold and harsh climatic conditions. Also, it remains unclear whether CsubMADS1 participates in other nutrient-limiting or excess environments, for example, phosphorus and sulphur. Furthermore, information about the cross-talk between chlorophyll A/B binding proteins and nitrogen remains scarce; hence more work needs to be done in this area. Therefore, this study has raised further questions regarding the potential roles of this transcription factor from *Coccomyxa subellipsoidea* C-169, which will hopefully be addressed in future work.

## 4. Experimental Procedures

### 4.1 Microalgal growth conditions and culture

*Coccomyxa subellipsoidea* C-169 WT cultures (accession number: NIES-2166) were obtained from the Microbial culture collection at the National Institute for Environmental Studies, Tsukuba, Japan. The *CsubMADS1* overexpressors and C-169 WT cultures were grown according to the method described in a previous study (Nayar and Thangavel, 2021). The stationary phase cultures were used for experiments.

### 4.2 Nitrogen Starvation

The nitrogen starvation protocol was followed according to an earlier study on *Chlamydomonas* with slight modifications (Kwak *et al*., 2017). For *Coccomyxa subellipsoidea* C-169, the cells (WT and *MADS* OX cells) were first inoculated into Modified Bold’s Basal medium (MBBM) for two days. Next, pre-cultures were initiated with the same cell density and were grown for three days. The cells were washed with fresh MBBM or MBBM-N (nitrogen deficient) media, and cells were pelleted by centrifugation at 4000 rpm for 3 mins. The cells were then inoculated in MBBM or MBBM-N with cell density 5 × 10^6^ and cultured for seven days. The cells were harvested on the seventh day for further experiments.

### 4.3 Staining lipid droplets with Nile Red and Confocal Microscopy

Stationary phase cultures and nitrogen starved (WT, OX1 and OX5) cultures were stained with Nile Red (Sigma-Aldrich, USA), as described in a previous study with minor modifications (Storms *et al*., 2014). Once the Nile red was added to the cell suspension in a microcentrifuge tube, the mixture was shaken at 200 rpm, 40 degrees, for 10 mins. The stained cells were centrifuged at 3600 rpm for 5 min and resuspended in 1X PBS, pH 7.6. A 10 µl aliquot was placed on a glass slide along with a coverslip and observed under the confocal microscope with an excitation laser 488 nm (Nikon, Japan). The bright field as well as fluorescence images were taken separately and overlaid by using Adobe R Photoshop CC.

### 4.4 Flow cytometric Analysis of Neutral lipids

Cells from stationary phase and nitrogen-starved cultures stained with Nile red were subjected to flow cytometric analysis in a BD FACS Aria™ III machine (BD Biosciences, USA) to analyze neutral lipids as described in an earlier study (Satpati and Pal, 2015).

### 4.5 Methylene Staining of cells

Methylene staining of stationary phase cells and nitrogen-starved cells (WT, OX1 and OX5) was done as described in an earlier study (Nayar and Thangavel, 2021).

### 4.6 RNA Isolation

Total RNA was isolated from the stationary phase, and nitrogen starvation treated WT, *CsubMADS1* OX1, and OX5 cultures using TRIzol reagent (Invitrogen) for three biological replicates, according to the manufacturer’s protocol (Invitrogen) as described in a previous study (Nayar and Thangavel, 2021).

### 4.7 RNA Sequencing

Total RNA was extracted using Trizol (Invitrogen, United States) from *Coccomyxa* cells (WT and *CsubMADS1* OX1, three biological replicates each) according to the manufacturer’s protocol. RNA sequencing was carried out as described in a previous study (Nayar, 2021). The *Coccomyxa subellipsoidea* C-169 reference genome was downloaded from the NCBI database. Because the aim of the RNA-Seq experiment was to obtain a shortlist of candidate target genes rather than to perform a comprehensive differential expression analysis, a simple fold-change and *p*-value–based approach was used to highlight genes showing consistent expression shifts across biological replicates. Normalized expression values obtained from DESeq were used to calculate fold changes for a given gene. The p-value for the individual genes was calculated using the Student’s two-tailed, unpaired t-test, with p < 0.05 was considered significant. Genes were categorized up- and downregulated based on the fold change cut-off of 1.5 together with p < 0.05. The RNA-Seq, raw and processed files have been deposited to the GEO database, NCBI, with accession number GSE214369.

### 4.8 Real-time PCR

The cDNA from the stationary phase and nitrogen-depleted cultures (WT, OX1, OX5; three biological replicates each) for qPCR was prepared with the total RNA using the cDNA archive kit (Thermo Fisher Scientific) as per the manufacturer’s protocol, as described earlier (Nayar and Thangavel, 2021). qPCR primers are listed in Table S6.

### 4.9 CsubMADS1 Protein expression and purification

The coding region (CDS) of *CsubMADS1* was amplified from *Coccomyxa subellipsoidea* cDNA. The primers used for the PCR were MADS pET28 F and MADS pET28 R (see Table S6). The cloning in the pET28a vector (Novagen), expression, and purification of CsubMADS1 protein in BL21 DE3 *E. coli* cells were done as described earlier (Nayar, 2021).

### 4.10 Western Blotting

Purified protein, wash flow through, and lysate flow through were denatured in Laemelli buffer (0.5 M Tris-HCl, pH 6.8, glycerol, 10% SDS, 0.5% bromophenol blue, 0.71 M β-mercaptoethanol) at 95°C for 5 mins. A 12% polyacrylamide gel (Bio-rad, USA) was loaded with the denatured protein samples and a protein marker (Bio-rad, USA) and electrophoresed at a constant voltage (200 V) in a mini protean tetra cell apparatus (Bio-rad, USA). Western blotting was carried out using antibodies Anti-CsubMADS1 (1:2500 dilution) (generated by using a small CsubMADS1 specific peptide from the MADS domain; RIEKIGDERNRQVTFTKRKN; Abgenex, Bhuvaneshwar) and Anti-His (Sigma-Aldrich, USA) (1:3000 dilution) as described in a previous study (Nayar *et al*., 2013).

### 4.11 Electromobility shift assays (EMSA)

Electromobility shift assays were carried out using 500 ng, 1000 ng, 1500 ng and 2000 ng of purified protein and 40 ng DNA probes containing the CArG boxes of *CAB24* and *CAB8* in separate reactions (see Table S6). Supershift assays were done using anti-CsubMADS1 and anti-His antibodies. These antibodies were already tested using western blotting for specificity. For all the assays, the Electrophoretic Mobility Shift Assay (EMSA) Kit from Molecular Probes (Invitrogen, USA) was used according to the manufacturer’s protocol. A 1.5% Agarose gel was used for all the assays. Agarose-based EMSA was used instead of polyacrylamide because preliminary assays indicated that CsubMADS1 formed higher-order DNA–protein complexes that were not clearly resolved on PAGE as it remained stuck in the wells. Agarose gels have been used successfully in EMSA when proteins produce large or multimeric complexes that are difficult to separate on polyacrylamide. The 1.5% agarose matrix allowed improved visualization of retarded complexes for CsubMADS1–CArG probes. These assays were repeated thrice. However, we acknowledge that PAGE remains the gold standard for EMSA and future validation with PAGE-based assays would strengthen these findings.

### 4.12 Statistics

For each sample, three biological replicates were included in each experiment. The two-tailed unpaired Student t-test was used to determine the results’ statistical significance (P < 0.05).

## Acknowledgements

SN sincerely thanks the SERB, Department of Science and Technology (DST), Government of India (GOI) for the Core Research Grant (File No: EMR/2016/001497), and DST, GOI for the INSPIRE Faculty Award. SN also sincerely thanks Rajiv Gandhi Centre for Biotechnology (RGCB), Thiruvananthapuram for hosting the INSPIRE Faculty Award and providing all necessary infrastructure. SN thanks the Central Instrumentation Facility team of RGCB. SN also thanks Genotypic Pvt. Ltd, Bengaluru, for performing the RNA Seq and initial data analysis for the data presented here.

## Author Contributions

SN acquired the funding, planned the work, performed the experiments, and wrote the manuscript.

## Supporting information

Figure S1- Wild type and overexpressors (OX1, OX5 and OX7) culture grown in nitrogen-deficient medium

Figure S2- Position of CArG boxes in the upstream region of *CAB* genes

Figure S3- SDS PAGE of induced lysate, lysate flow through, wash flow through and purified protein fraction.

Table S1- Top pathway upregulated in *CsubMADS1* OX

Table S2- Top Pathway downregulated in *CsubMADS1* OX

Table S3- List of *CAB*s upregulated in *CsubMADS1* OX

Table S4- List of CABs in *Coccomyxa subellipsoidea* C-169

Table S5- CArG boxes in the 2 kb upstream region of *CAB*s

Table S6- Primers and probes used in this study

## Declaration of generative AI and AI-assisted technologies in the manuscript preparation process

During the preparation of this work the author used ChatGPT for language editing of selected parts of the manuscript. The author reviewed and edited the content as needed and takes full responsibility for the content of the published article.

## References

Andersson, U., Heddad, M. and Adamska, I. (2003) Light stress-induced one-helix protein of the chlorophyll a/b-binding family associated with photosystem I. Plant Physiol, 132, 811–820.

Arora, R., Agarwal, P., Ray, S., Singh, A.K., Singh, V.P., Tyagi, A.K. and Kapoor, S. (2007) MADS-box gene family in rice: genome-wide identification, organization and expression profiling during reproductive development and stress. BMC Genomics, 8, 242.

Baek, D., Chun, H.J., Yun, D.J. and Kim, M.C. (2017) Cross-talk between Phosphate Starvation and Other Environmental Stress Signaling Pathways in Plants. Mol Cells, 40, 697–705.

Bajhaiya, A.K., Dean, A.P., Zeef, L.A., Webster, R.E. and Pittman, J.K. (2016) PSR1 Is a Global Transcriptional Regulator of Phosphorus Deficiency Responses and Carbon Storage Metabolism in Chlamydomonas reinhardtii. Plant Physiol, 170, 1216–1234.

Barragán-Rosillo, A.C., Peralta-Alvarez, C.A., Ojeda-Rivera, J.O., Arzate-Mejía, R.G., Recillas-Targa, F. and Herrera-Estrella, L. (2021) Genome accessibility dynamics in response to phosphate limitation is controlled by the PHR1 family of transcription factors in Arabidopsis. Proc Natl Acad Sci U S A, 118.

Bechtold, U., Penfold, C.A., Jenkins, D.J., Legaie, R., Moore, J.D., Lawson, T., Matthews, J.S., Vialet-Chabrand, S.R., Baxter, L., Subramaniam, S., Hickman, R., Florance, H., Sambles, C., Salmon, D.L., Feil, R., Bowden, L., Hill, C., Baker, N.R., Lunn, J.E., Finkenstädt, B., Mead, A., Buchanan-Wollaston, V., Beynon, J., Rand, D.A., Wild, D.L., Denby, K.J., Ott, S., Smirnoff, N. and Mullineaux, P.M. (2016) Time-Series Transcriptomics Reveals That AGAMOUS-LIKE22 Affects Primary Metabolism and Developmental Processes in Drought-Stressed Arabidopsis. Plant Cell, 28, 345–366.

Blanc, G., Agarkova, I., Grimwood, J., Kuo, A., Brueggeman, A., Dunigan, D.D., Gurnon, J., Ladunga, I., Lindquist, E., Lucas, S., Pangilinan, J., Proschold, T., Salamov, A., Schmutz, J., Weeks, D., Yamada, T., Lomsadze, A., Borodovsky, M., Claverie, J.M., Grigoriev, I.V. and Van Etten, J.L. (2012) The genome of the polar eukaryotic microalga Coccomyxa subellipsoidea reveals traits of cold adaptation. Genome Biol, 13, R39.

Cai, X., Gao, L., Teng, L., Ge, J., Oo, Z.M., Kumar, A.R., Gilliland, D.G., Mason, P.J., Tan, K. and Speck, N.A. (2015) Runx1 Deficiency Decreases Ribosome Biogenesis and Confers Stress Resistance to Hematopoietic Stem and Progenitor Cells. Cell Stem Cell, 17, 165–177.

Castelan-Munoz, N., Herrera, J., Cajero-Sanchez, W., Arrizubieta, M., Trejo, C., Garcia-Ponce, B., Sanchez, M.P., Alvarez-Buylla, E.R. and Garay-Arroyo, A. (2019) MADS-Box Genes Are Key Components of Genetic Regulatory Networks Involved in Abiotic Stress and Plastic Developmental Responses in Plants. Front Plant Sci, 10, 853.

Cavonius, L., Fink, H., Kiskis, J., Albers, E., Undeland, I. and Enejder, A. (2015) Imaging of lipids in microalgae with coherent anti-stokes Raman scattering microscopy. Plant Physiol, 167, 603–616.

Chen, C., Begcy, K., Liu, K., Folsom, J.J., Wang, Z., Zhang, C. and Walia, H. (2016) Heat stress yields a unique MADS box transcription factor in determining seed size and thermal sensitivity. Plant Physiol, 171, 606–622.

Chen, L., Zhao, Y., Xu, S., Zhang, Z., Xu, Y., Zhang, J. and Chong, K. (2018) OsMADS57 together with OsTB1 coordinates transcription of its target OsWRKY94 and D14 to switch its organogenesis to defense for cold adaptation in rice. New Phytol, 218, 219–231.

Chen, R., Ma, J., Luo, D., Hou, X., Ma, F., Zhang, Y., Meng, Y., Zhang, H. and Guo, W. (2019) CaMADS, a MADS-box transcription factor from pepper, plays an important role in the response to cold, salt, and osmotic stress. Plant Sci, 280, 164–174.

Cordeiro, T.N., Schmidt, H., Madrid, C., Juarez, A., Bernado, P., Griesinger, C., Garcia, J. and Pons, M. (2011) Indirect DNA readout by an H-NS related protein: structure of the DNA complex of the C-terminal domain of Ler. PLoS Pathog, 7, e1002380.

Curcuraci, E., Manuguerra, S., Messina, C.M., Arena, R., Renda, G., Ioannou, T., Amato, V., Hellio, C., Barba, F.J. and Santulli, A. (2022) Culture Conditions Affect Antioxidant Production, Metabolism and Related Biomarkers of the Microalgae Phaeodactylum tricornutum. Antioxidants (Basel), 11.

Das, D., Paries, M., Hobecker, K., Gigl, M., Dawid, C., Lam, H.M., Zhang, J., Chen, M. and Gutjahr, C. (2022) PHOSPHATE STARVATION RESPONSE transcription factors enable arbuscular mycorrhiza symbiosis. Nat Commun, 13, 477.

Deng, X., Yang, J., Wu, X., Li, Y. and Fei, X. (2014) A C2H2 zinc finger protein FEMU2 is required for fox1 expression in Chlamydomonas reinhardtii. PLoS One, 9, e112977.

Devaiah, B.N., Karthikeyan, A.S. and Raghothama, K.G. (2007) WRKY75 transcription factor is a modulator of phosphate acquisition and root development in Arabidopsis. Plant Physiol, 143, 1789–1801.

Dittami, S.M., Michel, G., Collen, J., Boyen, C. and Tonon, T. (2010) Chlorophyll-binding proteins revisited--a multigenic family of light-harvesting and stress proteins from a brown algal perspective. BMC Evol Biol, 10, 365.

Dittami, S.M., Scornet, D., Petit, J.-L., Ségurens, B., Da Silva, C., Corre, E., Dondrup, M., Glatting, K.-H., König, R., Sterck, L., Rouzé, P., Van de Peer, Y., Cock, J.M., Boyen, C. and Tonon, T. (2009) Global expression analysis of the brown alga Ectocarpus siliculosus (Phaeophyceae) reveals large-scale reprogramming of the transcriptome in response to abiotic stress. Genome Biology, 10, R66.

Dolan, J.W. and Fields, S. (1991) Cell-type-specific transcription in yeast. Biochim Biophys Acta, 1088, 155–169.

Fan, J., Cui, Y., Wan, M., Wang, W. and Li, Y. (2014) Lipid accumulation and biosynthesis genes response of the oleaginous Chlorella pyrenoidosa under three nutrition stressors.Biotechnol Biofuels, 7, 17.

Gan, Y., Bernreiter, A., Filleur, S., Abram, B. and Forde, B.G. (2012) Overexpressing the ANR1 MADS-box gene in transgenic plants provides new insights into its role in the nitrate regulation of root development. Plant Cell Physiol, 53, 1003–1016.

Gan, Y., Filleur, S., Rahman, A., Gotensparre, S. and Forde, B.G. (2005) Nutritional regulation of ANR1 and other root-expressed MADS-box genes in Arabidopsis thaliana. Planta, 222, 730–742.

Goncalves, E.C., Koh, J., Zhu, N., Yoo, M.J., Chen, S., Matsuo, T., Johnson, J.V. and Rathinasabapathi, B. (2016) Nitrogen starvation-induced accumulation of triacylglycerol in the green algae: evidence for a role for ROC40, a transcription factor involved in circadian rhythm. Plant J, 85, 743–757.

Guan, P., Ripoll, J.J., Wang, R., Vuong, L., Bailey-Steinitz, L.J., Ye, D. and Crawford, N.M. (2017) Interacting TCP and NLP transcription factors control plant responses to nitrate availability. Proc Natl Acad Sci U S A, 114, 2419–2424.

Guo, X., Chen, G., Cui, B., Gao, Q., Guo, J.-E., Li, A., Zhang, L. and Hu, Z. (2016) Solanum lycopersicum agamous-like MADS-box protein AGL15-like gene, SlMBP11, confers salt stress tolerance. Molecular Breeding, 36, 125.

Hamill, P.G., Stevenson, A., McMullan, P.E., Williams, J.P., Lewis, A.D.R., S, S., Stevenson, K.E., Farnsworth, K.D., Khroustalyova, G., Takemoto, J.Y., Quinn, J.P., Rapoport, A. and Hallsworth, J.E. (2020) Microbial lag phase can be indicative of, or independent from, cellular stress. Sci Rep, 10, 5948.

Hanakahi, L.A., Dempsey, L.A., Li, M.J. and Maizels, N. (1997) Nucleolin is one component of the B cell-specific transcription factor and switch region binding protein, LR1. Proc Natl Acad Sci U S A, 94, 3605–3610.

He, Q., Dolganov, N., Bjorkman, O. and Grossman, A.R. (2001) The high light-inducible polypeptides in Synechocystis PCC6803. Expression and function in high light. J Biol Chem, 276, 306–314.

Hidayati, N.A., Yamada-Oshima, Y., Iwai, M., Yamano, T., Kajikawa, M., Sakurai, N., Suda, K., Sesoko, K., Hori, K., Obayashi, T., Shimojima, M., Fukuzawa, H. and Ohta, H. (2019) Lipid remodeling regulator 1 (LRL1) is differently involved in the phosphorus-depletion response from PSR1 in Chlamydomonas reinhardtii. Plant J, 100, 610–626.

Hwang, Y.S., Jung, G. and Jin, E. (2008) Transcriptome analysis of acclimatory responses to thermal stress in Antarctic algae. Biochem Biophys Res Commun, 367, 635–641.

Imamura, S., Kanesaki, Y., Ohnuma, M., Inouye, T., Sekine, Y., Fujiwara, T., Kuroiwa, T. and Tanaka, K. (2009) R2R3-type MYB transcription factor, CmMYB1, is a central nitrogen assimilation regulator in Cyanidioschyzon merolae. Proc Natl Acad Sci U S A, 106, 12548–12553.

Jing, D., Agnew, J., Patton, W.F., Hendrickson, J. and Beechem, J.M. (2003) A sensitive two-color electrophoretic mobility shift assay for detecting both nucleic acids and protein in gels. Proteomics, 3, 1172–1180.

Kang, N.K., Jeon, S., Kwon, S., Koh, H.G., Shin, S.E., Lee, B., Choi, G.G., Yang, J.W., Jeong, B.R. and Chang, Y.K. (2015) Effects of overexpression of a bHLH transcription factor on biomass and lipid production in Nannochloropsis salina. Biotechnol Biofuels, 8, 200.

Kater, M.M., Dreni, L. and Colombo, L. (2006) Functional conservation of MADS-box factors controlling floral organ identity in rice and Arabidopsis. J Exp Bot, 57, 3433–3444.

Khong, G.N., Pati, P.K., Richaud, F., Parizot, B., Bidzinski, P., Mai, C.D., Bes, M., Bourrie, I., Meynard, D., Beeckman, T., Selvaraj, M.G., Manabu, I., Genga, A.M., Brugidou, C., Nang Do, V., Guiderdoni, E., Morel, J.B. and Gantet, P. (2015) OsMADS26 Negatively Regulates Resistance to Pathogens and Drought Tolerance in Rice. Plant Physiol, 169, 2935–2949.

Koshimizu, S., Kofuji, R., Sasaki-Sekimoto, Y., Kikkawa, M., Shimojima, M., Ohta, H., Shigenobu, S., Kabeya, Y., Hiwatashi, Y., Tamada, Y., Murata, T. and Hasebe, M. (2018) Physcomitrella MADS-box genes regulate water supply and sperm movement for fertilization. Nat Plants, 4, 36–45.

Krapp, A., Berthomé, R., Orsel, M., Mercey-Boutet, S., Yu, A., Castaings, L., Elftieh, S., Major, H., Renou, J.-P. and Daniel-Vedele, F. (2011) Arabidopsis Roots and Shoots Show Distinct Temporal Adaptation Patterns toward Nitrogen Starvation Plant Physiology, 157, 1255–1282.

Kumar Sharma, A., Mühlroth, A., Jouhet, J., Maréchal, E., Alipanah, L., Kissen, R., Brembu, T., Bones, A.M. and Winge, P. (2020) The Myb-like transcription factor phosphorus starvation response (PtPSR) controls conditional P acquisition and remodelling in marine microalgae. New Phytol, 225, 2380–2395.

Kwak, M., Park, W.-K., Shin, S.-E., Koh, H.-G., Lee, B., Jeong, B.-r. and Chang, Y.K. (2017) Improvement of biomass and lipid yield under stress conditions by using diploid strains of Chlamydomonas reinhardtii. Algal Research, 26, 180–189.

Lambertz, C., Hemschemeier, A. and Happe, T. (2010) Anaerobic expression of the ferredoxin-encoding FDX5 gene of Chlamydomonas reinhardtii is regulated by the Crr1 transcription factor. Eukaryot Cell, 9, 1747–1754.

Lee, J.H., Yoo, S.J., Park, S.H., Hwang, I., Lee, J.S. and Ahn, J.H. (2007) Role of SVP in the control of flowering time by ambient temperature in Arabidopsis. Genes Dev, 21, 397–402.

Leseberg, C.H., Li, A., Kang, H., Duvall, M. and Mao, L. (2006) Genome-wide analysis of the MADS-box gene family in Populus trichocarpa. Gene, 378, 84–94.

Li, D., Liu, C., Shen, L., Wu, Y., Chen, H., Robertson, M., Helliwell, C.A., Ito, T., Meyerowitz, E. and Yu, H. (2008) A repressor complex governs the integration of flowering signals in Arabidopsis. Dev Cell, 15, 110–120.

Li, X.W., Zhu, Y.L., Chen, C.Y., Geng, Z.J., Li, X.Y., Ye, T.T., Mao, X.N. and Du, F. (2020) Cloning and characterization of two chlorophyll A/B binding protein genes and analysis of their gene family in Camellia sinensis. Sci Rep, 10, 4602.

Liu, R., Xu, Y.H., Jiang, S.C., Lu, K., Lu, Y.F., Feng, X.J., Wu, Z., Liang, S., Yu, Y.T., Wang, X.F. and Zhang, D.P. (2013) Light-harvesting chlorophyll a/b-binding proteins, positively involved in abscisic acid signalling, require a transcription repressor, WRKY40, to balance their function. J Exp Bot, 64, 5443–5456.

Luang, S., Sornaraj, P., Bazanova, N., Jia, W., Eini, O., Hussain, S.S., Kovalchuk, N., Agarwal, P.K., Hrmova, M. and Lopato, S. (2018) The wheat TabZIP2 transcription factor is activated by the nutrient starvation-responsive SnRK3/CIPK protein kinase. Plant Mol Biol, 96, 543–561.

Matthijs, M., Fabris, M., Obata, T., Foubert, I., Franco-Zorrilla, J.M., Solano, R., Fernie, A.R., Vyverman, W. and Goossens, A. (2017) The transcription factor bZIP14 regulates the TCA cycle in the diatom Phaeodactylum tricornutum. Embo j, 36, 1559–1576.

Millar, A.J. and Kay, S.A. (1996) Integration of circadian and phototransduction pathways in the network controlling CAB gene transcription in Arabidopsis. Proc Natl Acad Sci U S A, 93, 15491–15496.

Moin, M., Saha, A., Bakshi, A., Madhav, M.S. and Kirti, P.B. (2020) Ribosomal Protein Large subunit RPL6 modulates salt tolerance in rice. bioRxiv, 2020.2005.2031.126102.

Møller, A.L., Pedas, P., Andersen, B., Svensson, B., Schjoerring, J.K. and Finnie, C. (2011) Responses of barley root and shoot proteomes to long-term nitrogen deficiency, short-term nitrogen starvation and ammonium. Plant Cell Environ, 34, 2024–2037.

Msanne, J., Xu, D., Konda, A.R., Casas-Mollano, J.A., Awada, T., Cahoon, E.B. and Cerutti, H. (2012) Metabolic and gene expression changes triggered by nitrogen deprivation in the photoautotrophically grown microalgae Chlamydomonas reinhardtii and Coccomyxa sp. C-169. Phytochemistry, 75, 50–59.

Murakami, H., Kakutani, N., Kuroyanagi, Y., Iwai, M., Hori, K., Shimojima, M. and Ohta, H. (2020) MYB-like transcription factor NoPSR1 is crucial for membrane lipid remodeling under phosphate starvation in the oleaginous microalga Nannochloropsis oceanica. FEBS Lett, 594, 3384–3394.

Nagaraj, S., Senthil-Kumar, M., Ramu, V.S., Wang, K. and Mysore, K.S. (2016) Plant Ribosomal Proteins, RPL12 and RPL19, Play a Role in Nonhost Disease Resistance against Bacterial Pathogens. Frontiers in Plant Science, 6.

Nayar, S. (2021) Exploring the Role of a Cytokinin-Activating Enzyme LONELY GUY in Unicellular Microalga Chlorella variabilis. Front Plant Sci, 11, 611871.

Nayar, S., Sharma, R., Tyagi, A.K. and Kapoor, S. (2013) Functional delineation of rice MADS29 reveals its role in embryo and endosperm development by affecting hormone homeostasis. J Exp Bot, 64, 4239–4253.

Nayar, S. and Thangavel, G. (2021) CsubMADS1, a lag phase transcription factor, controls development of polar eukaryotic microalga Coccomyxa subellipsoidea C-169. Plant J, 107, 1228–1242.

Parenicová, L., de Folter, S., Kieffer, M., Horner, D.S., Favalli, C., Busscher, J., Cook, H.E., Ingram, R.M., Kater, M.M., Davies, B., Angenent, G.C. and Colombo, L. (2003) Molecular and phylogenetic analyses of the complete MADS-box transcription factor family in Arabidopsis: new openings to the MADS world. Plant Cell, 15, 1538–1551.

Park, S., Jung, G., Hwang, Y.S. and Jin, E. (2010) Dynamic response of the transcriptome of a psychrophilic diatom, Chaetoceros neogracile, to high irradiance. Planta, 231, 349–360.

Riboni, M., Galbiati, M., Tonelli, C. and Conti, L. (2013) GIGANTEA enables drought escape response via abscisic acid-dependent activation of the florigens and SUPPRESSOR OF OVEREXPRESSION OF CONSTANS. Plant Physiol, 162, 1706–1719.

Riboni, M., Robustelli Test, A., Galbiati, M., Tonelli, C. and Conti, L. (2016) ABA-dependent control of GIGANTEA signalling enables drought escape via up-regulation of FLOWERING LOCUS T in Arabidopsis thaliana. J Exp Bot, 67, 6309–6322.

Ried, M.K., Wild, R., Zhu, J., Pipercevic, J., Sturm, K., Broger, L., Harmel, R.K., Abriata, L.A., Hothorn, L.A., Fiedler, D., Hiller, S. and Hothorn, M. (2021) Inositol pyrophosphates promote the interaction of SPX domains with the coiled-coil motif of PHR transcription factors to regulate plant phosphate homeostasis. Nat Commun, 12, 384.

Satpati, G.G. and Pal, R. (2015) Rapid detection of neutral lipid in green microalgae by flow cytometry in combination with Nile red staining—an improved technique. Annals of Microbiology, 65, 937–949.

Shibata, M., Favero, D.S., Takebayashi, R., Takebayashi, A., Kawamura, A., Rymen, B., Hosokawa, Y. and Sugimoto, K. (2022) Trihelix transcription factors GTL1 and DF1 prevent aberrant root hair formation in an excess nutrient condition. New Phytol, 235, 1426–1441.

Shiraku, M.L., Magwanga, R.O., Cai, X., Kirungu, J.N., Xu, Y., Mehari, T.G., Hou, Y., Wang, Y., Wang, K., Peng, R., Zhou, Z. and Liu, F. (2021) Knockdown of 60S ribosomal protein L14-2 reveals their potential regulatory roles to enhance drought and salt tolerance in cotton. Journal of Cotton Research, 4, 27.

Silverthorne, J. and Tobin, E.M. (1984) Demonstration of transcriptional regulation of specific genes by phytochrome action. Proc Natl Acad Sci U S A, 81, 1112–1116.

Sommer, F., Kropat, J., Malasarn, D., Grossoehme, N.E., Chen, X., Giedroc, D.P. and Merchant, S.S. (2010) The CRR1 nutritional copper sensor in Chlamydomonas contains two distinct metal-responsive domains. Plant Cell, 22, 4098–4113.

Staneloni, R.J., Rodriguez-Batiller, M.J. and Casal, J.J. (2008) Abscisic acid, high-light, and oxidative stress down-regulate a photosynthetic gene via a promoter motif not involved in phytochrome-mediated transcriptional regulation. Mol Plant, 1, 75–83.

Storms, Z.J., Cameron, E., de la Hoz Siegler, H. and McCaffrey, W.C. (2014) A simple and rapid protocol for measuring neutral lipids in algal cells using fluorescence. J Vis Exp.

Tang, W. and Perry, S.E. (2003) Binding site selection for the plant MADS domain protein AGL15: an in vitro and in vivo study. J Biol Chem, 278, 28154–28159.

Thangavel, G. and Nayar, S. (2018) A Survey of MIKC Type MADS-Box Genes in Non-seed Plants: Algae, Bryophytes, Lycophytes and Ferns. Front Plant Sci, 9, 510.

Treisman, R. (1992) The serum response element. Trends Biochem Sci, 17, 423–426.

Valledor, L., Furuhashi, T., Hanak, A.M. and Weckwerth, W. (2013) Systemic cold stress adaptation of Chlamydomonas reinhardtii. Mol Cell Proteomics, 12, 2032–2047.

Varsano, T., Wolf, S.G. and Pick, U. (2006) A chlorophyll a/b-binding protein homolog that is induced by iron deficiency is associated with enlarged photosystem I units in the eucaryotic alga Dunaliella salina. J Biol Chem, 281, 10305–10315.

Wang, Z., Wang, F., Hong, Y., Yao, J., Ren, Z., Shi, H. and Zhu, J.-K. (2018) The Flowering Repressor SVP Confers Drought Resistance in Arabidopsis by Regulating Abscisic Acid Catabolism. Molecular Plant, 11, 1184–1197.

Warner, J.R. and McIntosh, K.B. (2009) How common are extraribosomal functions of ribosomal proteins? Mol Cell, 34, 3–11.

West, A.G., Shore, P. and Sharrocks, A.D. (1997) DNA binding by MADS-box transcription factors: a molecular mechanism for differential DNA bending. Mol Cell Biol, 17, 2876–2887.

Wilkinson, O.J., Martin-Gonzalez, A., Kang, H., Northall, S.J., Wigley, D.B., Moreno-Herrero, F. and Dillingham, M.S. (2019) CtIP forms a tetrameric dumbbell-shaped particle which bridges complex DNA end structures for double-strand break repair. Elife, 8.

Wool, I.G. (1996) Extraribosomal functions of ribosomal proteins. Trends Biochem Sci, 21, 164–165.

Wu, R., Liu, Z., Wang, J., Guo, C., Zhou, Y., Bawa, G., Rochaix, J.D. and Sun, X. (2022) COE2 Is Required for the Root Foraging Response to Nitrogen Limitation. Int J Mol Sci, 23.

Wykoff, D.D., Grossman, A.R., Weeks, D.P., Usuda, H. and Shimogawara, K. (1999) Psr1, a nuclear localized protein that regulates phosphorus metabolism in Chlamydomonas. Proc Natl Acad Sci U S A, 96, 15336–15341.

Xu, Y.H., Liu, R., Yan, L., Liu, Z.Q., Jiang, S.C., Shen, Y.Y., Wang, X.F. and Zhang, D.P. (2012) Light-harvesting chlorophyll a/b-binding proteins are required for stomatal response to abscisic acid in Arabidopsis. J Exp Bot, 63, 1095–1106.

Yang, T., Hao, L., Yao, S., Zhao, Y., Lu, W. and Xiao, K. (2016) TabHLH1, a bHLH-type transcription factor gene in wheat, improves plant tolerance to Pi and N deprivation via regulation of nutrient transporter gene transcription and ROS homeostasis. Plant Physiol Biochem, 104, 99–113.

Yu, C., Liu, Y., Zhang, A., Su, S., Yan, A., Huang, L., Ali, I., Liu, Y., Forde, B.G. and Gan, Y. (2015) MADS-box transcription factor OsMADS25 regulates root development through affection of nitrate accumulation in rice. PLoS One, 10, e0135196.

Yu, C., Su, S., Xu, Y., Zhao, Y., Yan, A., Huang, L., Ali, I. and Gan, Y. (2014) The effects of fluctuations in the nutrient supply on the expression of five members of the AGL17 clade of MADS-box genes in rice. PLoS One, 9, e105597.

Zhao, Y., Kong, H., Guo, Y. and Zou, Z. (2020) Light-harvesting chlorophyll a/b-binding protein-coding genes in jatropha and the comparison with castor, cassava and arabidopsis. PeerJ, 8, e8465.

Zobell, O., Faigl, W., Saedler, H. and Münster, T. (2010) MIKC* MADS-box proteins: conserved regulators of the gametophytic generation of land plants. Mol Biol Evol, 27, 1201–1211.

