## Supplementary figures and images for "CsubMADS1 overexpression is associated with increased CAB transcript levels and with enhanced tolerance under nutrient limitation"

### Figure S1

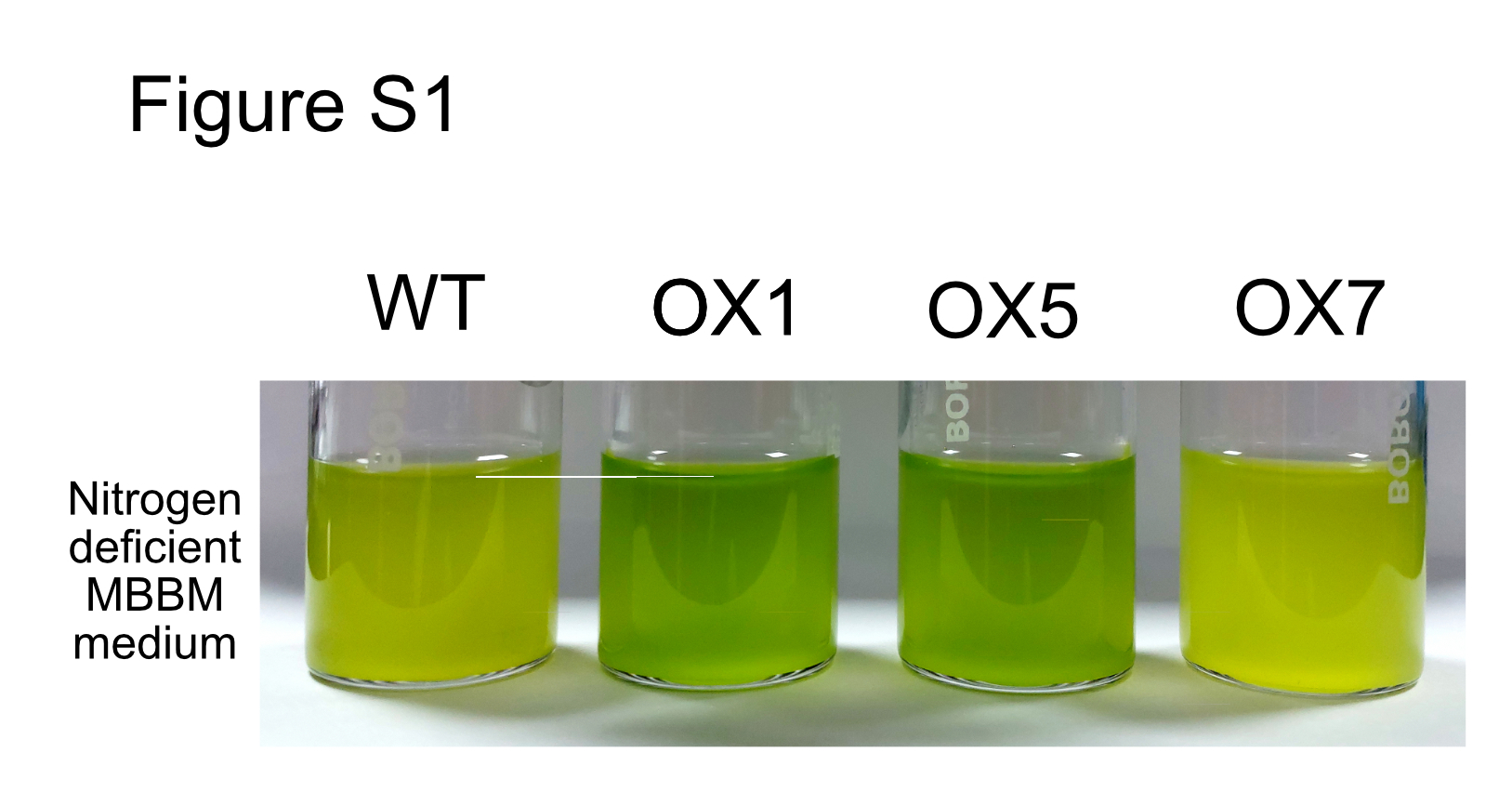

### Figure S2

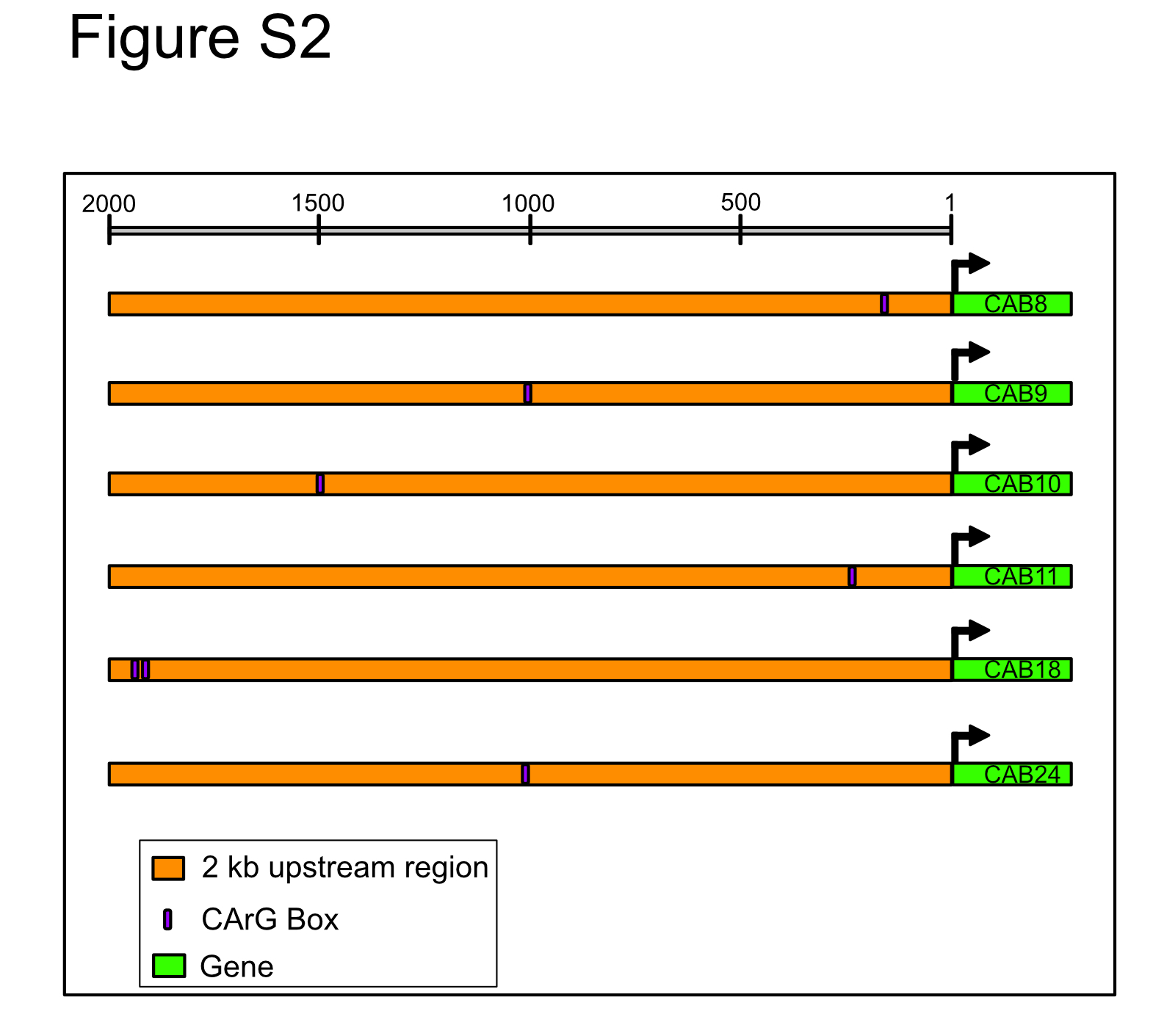

### Figure S3

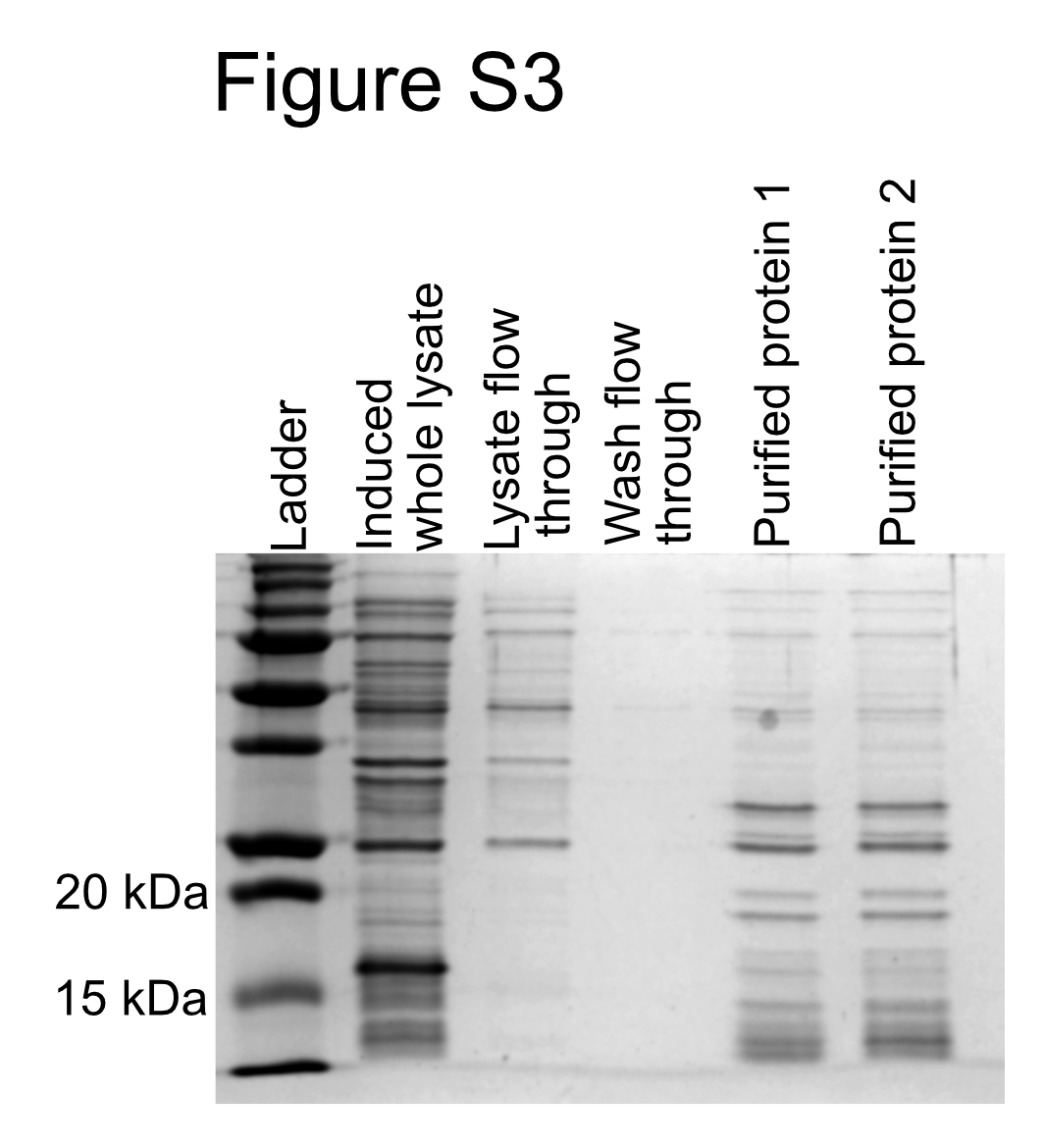
